# Intermittent fasting and caloric restriction interact with genetics to shape physiological health in mice

**DOI:** 10.1101/2021.04.02.438251

**Authors:** Guozhu Zhang, Andrew Deighan, Anil Raj, Laura Robinson, Hannah J. Donato, Gaven Garland, Mackenzie Leland, Baby Martin-McNulty, Ganesh A. Kolumam, Johannes Riegler, Adam Freund, Kevin M. Wright, Gary Churchill

**Affiliations:** Calico Life Sciences LLC, South San Francisco, California; The Jackson Laboratory, Bar Harbor, Maine

**Keywords:** intermittent fasting, caloric restriction, physiological health, gene x environ-ment interaction, Diversity Outcross mice

## Abstract

Dietary interventions can dramatically affect physiological health and organismal lifespan. The degree to which organismal health is improved depends upon genotype and the severity of dietary intervention, but neither the effects of these factors, nor their interaction, have been quantified in an outbred population. Moreover, it is not well understood what physiological changes occur shortly after dietary change and how these may affect the health of early adulthood population. In this article, we investigated the effect of six month exposure of either caloric restriction or intermittent fasting on a broad range of physiological traits in 960 one year old Diversity Outbred mice. We found caloric restriction and intermittent fasting affected distinct aspects of physiology and neither the magnitude nor the direction (beneficial or detrimental) of effects were concordant with the severity of the intervention. In addition to the effects of diet, genetic variation significantly affected 31 of 36 traits (heritabilties ranged from 0.04-0.65). We observed significant covariation between many traits that was due to both diet and genetics and quantified these effects with phenotypic and genetic correlations. We genetically mapped 16 diet-independent and 2 diet-dependent significant quantitative trait loci, both of which were associated with cardiac physiology. Collectively, these results demonstrate the degree to which diet and genetics interact to shape the physiological health of early adult-hood mice following six months of dietary intervention.

## Introduction

Dietary modifications are the most robust interventions known to increase organismal lifespan. Caloric restriction (CR) has been been shown to increase lifespan in multiple species including yeast, worms, flies, rats, mice, and non-human primates (Heilbronn and Ravussin (2003); Kaeberlein *et al*. (2005); Colman *et al*. (2009); Mattison *et al*. (2017); Liang *et al*. (2018); Pifferi *et al*. (2019)). Another dietary modification, intermittent fasting (IF), has been shown to increase lifespan in rodents (Goodrick *et al*. (1990)). However, the beneficial effects of these dietary interventions are not universal and can be influenced by sex, genetic variation and adaptation to the lab environment (Goodrick *et al*. (1990); Harper *et al*. (2006); Liao *et al*. (2010); Mitchell *et al*. (2016)). Moreover, the timing and duration of dietary intervention can alter the magnitude of lifespan effects, with the greatest increase observed when CR is imposed early and maintained throughout life (Weindruch *et al*. (1982); Yu *et al*. (1985); Goodrick *et al*. (1990); Dhahbi *et al*. (2004)). However, the age-specific genetic and physiological mechanisms that determine whether CR or IF will lengthen lifespan remain largely unknown.

Dietary intervention is hypothesized to extend lifespan by improving the physiological function of multiple systems, including but not limited to, metabolic, neurological, and cardiovascular (Ahmet *et al*. (2011); Colman *et al*. (2009); Gredilla and Barja (2005); Redman *et al*. (2018); Gräff *et al*. (2013); Patel *et al*. (2005); Halagappa *et al*. (2007)). In some instances, changes in gene expression, metabolite levels, and physiology occur shortly after the initiation of daily CR (Cao *et al*. (2001); Dhahbi *et al*. (2004); Mulligan *et al*. (2008); Bruss *et al*. (2010)). Despite the large number of CR experiments, it is not well understood how diet and genetics shape early-life changes in physiological traits and whether these changes may have lasting effects on lifespan.

In humans, the largest CR intervention trial published to date found that a two-year 25% CR treatment in a population of middle-aged, non-obese individuals caused significant reductions to multiple cardiovascular and metabolic syndrome risk factors (Kraus *et al*. (2019)). However, the effect of CR was not universally beneficial, participants in this trial experienced significant reductions in bone mineral density, muscle size and function (Villareal *et al*. (2006); Weiss *et al*. (2007); Villareal *et al*. (2016)). These studies demonstrate that CR improved multiple aspects of physiological function while worsening others in a relatively healthy population. It remains to be determined whether this result is a generalizable feature of CR interventions and whether IF treatment would produce similarly heterogeneous physiological effects. Additionally, it is unknown how genetic variation may contribute to the variation in the physiological response to dietary intervention.

We investigate the effect of both CR and IF on a range of physiological traits using Diversity Outbred (DO) mice (*Mus musculus*), a multi-parent genetic mapping population founded from eight in-bred strains (Svenson *et al*. (2012); Churchill *et al*. (2012)). Our goal is to identify how dietary interventions affect different aspects of physiology in early adulthood mice. We measure the effect of CR and IF on 36 morphological and functional traits derived from six phenotypic assays: grip strength, rotarod, dual-energy X-ray absorptiometry (DEXA), echocardiogram, acoustic startle, and wheel running. Many traits change significantly in one year old mice exposed to dietary intervention for six months. The correlated change in trait values enabled us to cluster traits into distinct axes of physiology and measure how they were altered in response to dietary intervention. A significant proportion of phenotypic variation in 30 traits is heritable and for many traits in the same cluster, a large proportion of the heritable variation the genetic effects are correlated. We map 24 diet-independent quantitative trait loci (QTL) and five diet-dependent QTLs. We impute all DO founder variants, fine-map QTL intervals to near single gene resolution and identify the founder allele(s) associated with trait variation. These findings enable us to conclude that dietary intervention has heterogeneous effects on physiological health in mice during early adulthood, phenotypic variation in many physiological health traits has a large genetic component, and in the case of cardiac physiology, variation is influenced by the interaction between genetics and dietary intervention.

## Study Design and Measurements

The DO mouse population was derived from eight inbred founder strains and is maintained at The Jackson Laboratory as an outbred heterozygous population (Svenson *et al*. (2012)). This study contains 960 female DO mice, sampled at generations: 22 – 24 and 26 – 28. There were two cohorts per generation for a total of 12 cohorts and 80 animals per cohort. Enrollment occurred in successive quarterly waves starting in March 2016 and continuing through November 2017.

A single female mouse per litter was enrolled into the study after wean age (three weeks old), so that no mice in the study were siblings and maximum genetic diversity was achieved. Mice were housed in pressurized, individually ventilated cages at a density of eight animals per cage (cage assignments were random). Mice were subject to a 12 hr:12 hr light:dark cycle beginning at 0600 hrs.

All animal procedures were approved by the Animal Care and Use Committee at The Jackson Laboratory.

From enrollment until six months of age, all mice were on an Ad Libitum diet of standard rodent chow 5KOG from LabDiet. At six months of age, each cage of eight animals was randomly assigned to one of five dietary treatments, with each cohort equally split between the five groups (N=192/group): Ad Libitum (AL), 20% caloric restriction (20), 40% caloric restriction (40), one day per week fast, (1D) and two days per week fast (2D) (see Figure 1). In a previous internal study at The Jackson Laboratory, the average food consumption of female DO mice was estimated to be 3.43g/day. Based on this observation, mice on 20% CR diet were given 2.75g/mouse/day and those on 40% CR diet were given 2.06g/mouse/day. Food was weighed out for an entire cage of eight. Observation of the animals indicated that the distribution of food was roughly equal among all mice in a cage across diet groups.

**Fig. 1.**
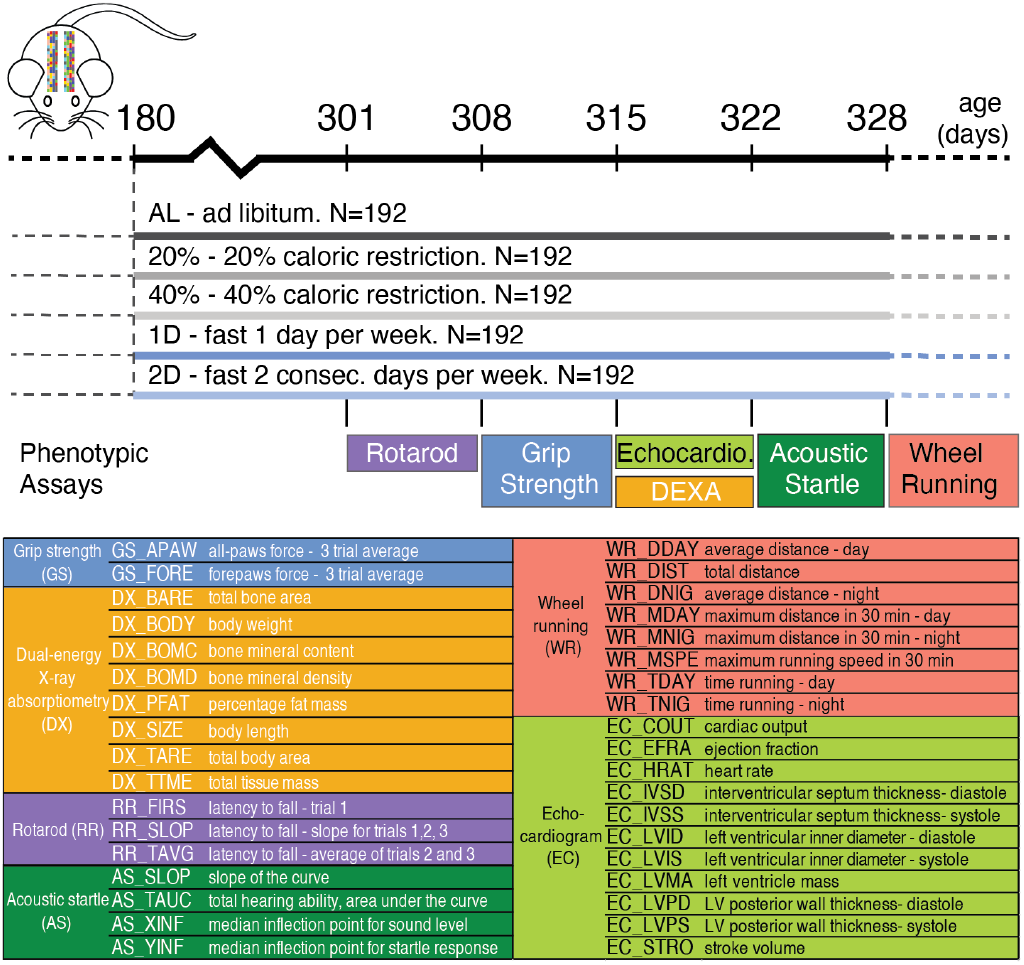
Study design. Dietary intervention starts at 180 days of age. Experimental procedures take approximately one week starting from given day.

Mice on AL diet had unlimited food access; they were fed when the cage was changed once a week. In rare instances when the AL mice consumed all food before the end of the week, the food was topped off mid week. Mice on 20% and 40% CR diets were fed daily. These mice were given a triple feeding on Friday afternoon to last till Monday afternoon. As the number of these mice in each cage decreased over time, the amount of food given to each cage was adjusted to reflect the number of mice in that cage. Fasting was imposed weekly from Wednesday noon to Thursday noon for mice on 1D diet and Wednesday noon to Friday noon for mice on 2D diet. Mice on 1D and 2D diets have unlimited food access (similar to AL mice) on their non-fasting days.

### Phenotypic Assays

We carried out six phenotypic assays to assess motor and neuromuscular function, activity, body composition, hearing and cardiovascular physiology, at approximately one year of age, five to six months following dietary intervention (Figure 1). All assays were conducted at The Jackson Laboratory following standard operating procedures that are included in the Supplemental Materials.

The rotarod assay was run with three consecutive trials per animal and we derived three traits to measure each animal’s latency to fall (Figure 1). The grip strength assay was run with three consecutive trials for all-paws and three trials for forepaws. In order to maximize the robustness of this assay, we removed any trial with log-normal Euclidean distance in the upper 5% quantile of the distribution of all animals and then calculated the per mouse average of the remaining trials. We used dual-energy X-ray absorptiometry to quantify eight body and bone composition traits (Figure 1). We measured voluntary wheel running in 30 minutes intervals for three nights and two days (mice were single housed for this assay). We used these data to derive average distance, time spent running and max speed in the following intervals: 12 hour day, 12 hour night and 24 hour intervals (Figure 1). The echocardiogram assay measured 11 traits capturing both heart morphology and function (Figure 1). Note, cardiac output is not directly measured, it is calculated from the product of stroke volume and heart rate.

The acoustic startle assay followed the sound-startle response protocol in which animals were exposed to five sound levels ranging from 80-120 decibels(dB) at 10dB steps. Each animal’s average startle response was normalized to background noise. To robustly measure hearing and sensorimotor function, we fit the startle response measurements for each animal to a five parameter logistic model with the R package nplr (Commo and Bot (2016)) and derived four values to quantify the shape of the logistic model (description provided in Figure 1). For a few animals, we estimated the the median sound response value to be greater than 120dB, the maximum sound level in our experiment. These values were set to 122dB, which is twice as loud as 120dB and is often used as the peak sound level in noise induced hearing loss research in rodent models (Kim *et al*. (2005); Escabi *et al*. (2019)).

### Outlier detection and batch correction

We first identified technical outliers resulting from equipment failure or mislabeled animals and if we could not manually correct them using lab records, they were removed. The total number of samples per trait after outlier removal is listed in Supplemental Table S1. In order to prevent potential biases in interpretation and increase the reliability of these trait measurements, we corrected values for batch and technician effects (Mandillo *et al*. (2008); Gulinello *et al*. (2019); Kafkafi *et al*. (2018)). For this experiment, there were 12 batches (two for each DO generation) and eight technicians. To quantify batch and technician effects, we fit an Analysis of Variance (ANOVA) model as follows:

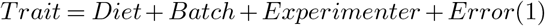

In contrast to all other assays, greater than 80% of echocardiogram derived traits were collected by a single technician and we determined that a reduced ANOVA model including Batch and not Experimenter terms was sufficient to control for the batch and technician effects. We used the residuals from each model to identify and remove biologically impossible values according to Tukey’s rule for far outlier (Tukey (1977)). After removing far outliers, we repeated the model fit procedure. To remove batch and experimenter effects, we adjusted each trait using the batch and experimenter model coefficients.

Grip strength and rotarod derived traits can be confounded by body weight (Crawley (2007); Maurissen *et al*. (2003); Hood (2011)) and in order to account for this, we fit the following Analysis of Covariance (ANCOVA) model:

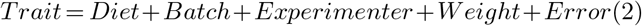

To remove body weight effect for grip strength and rotarod derived traits, we adjusted the trait value using the following formula:

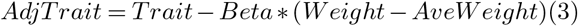

where *Beta* is the body weight coefficient from the ANCOVA model and *AveW eight* is the population mean body weight. Following technical, batch, technician, and outlier correction, we applied z-score standardization for all traits. Unless otherwise stated, these values were used for each subsequent analysis.

### Phenotypic effect of dietary intervention

In order to quantify the effect of dietary intervention on each trait, we applied an ANOVA model with Dunnett post-hoc test to compare each diet intervention group to the AL group. In order to account for statistical testing across multiple traits, we applied the Westfall-Young multiple testing adjustment (Westfall *et al*. (1993)).

### Phenotypic correlation and unsupervised clustering analysis

We calculated correlation coefficients within each diet treatment, and experiment-wide correlation coefficients for all animals across all diets. We performed unsupervised hierarchical clustering analysis using the distance metric 1 − |Phenotypic Correlation| and complete linkage. For animals to be included in this hierarchical clustering procedure we required they had no missing trait data (*N* = 525). To determine cluster membership of each trait, we applied a sensitivity analysis by first calculating the within cluster similarity, dist-within, of a trait as the average pairwise distance to all other traits in the same cluster. Second, we calculated the across cluster similarity, dist-across, as the average pairwise distance to all traits outside of the cluster. Small values of dist-within indicate the trait is highly correlated with traits within the cluster, whereas large values of dist-across indicate the trait is highly uncorrelated with traits outside of the cluster. To identify robust clusters of highly correlated traits, the hierarchical clustering algorithm minimizes the penalty score, defined as dist-within/dist-across. This penalty score is sensitive to the size of the cluster and to derive a cluster size specific penalty significance threshold, we used a bootstrap method with 1,000 resampling trials. The cluster size specific penalty significance threshold was defined as the 0.05/(cluster size-1) quantile value (Supplemental Table S2).

In order to organize traits into robust clusters, we first created a dendrogram with 5 clusters and compared the observed penalty scores to the bootstrap derived penalty threshold values. Within each cluster, we removed traits that had a higher penalty score than the penalty significance threshold by moving the cut-tree function closer to the origin node of the dendrogram. After the creation of a new set of clusters, we repeated the process until every newly created cluster had a penalty score that was less than the bootstrap derived penalty threshold values. We kept singletons, as single-trait clusters. Finally, we used principle component (PC) analysis of traits within the same multi-trait cluster to derive composite traits. All PC derived traits with a cumulative of 90% total variance explained were included in genetic linkage analyses.

### Genotype data and quality assessment

We collected tail clippings and extracted DNA using DNeasy Blood and Tissue Kit (Qiagen) from 954 animals. Samples were genotyped using the 143,259-probe GigaMUGA array from the Illumina Infinium II platform (Morgan *et al*. (2016)) by NeoGen Corp. (genomics.neogen.com/). We evaluated genotype quality using the R package: qtl2 (Broman *et al*. (2019)). We processed all raw genotype data with a corrected physical map of the GigaMUGA array probes (https://kbroman.org/MUGAarrays/muga_annotations.html). After filtering genetic markers for uniquely mapped probes, genotype quality and a 20% genotype missingness threshold, our dataset contained 110,807 markers.

We next examined the genotype quality of individual animals. We found seven pairs of animals with identical genotypes which suggested that one of each pair was mislabelled. We identified and removed a single mislabelled animal per pair by referencing the genetic data against coat color. Next, we removed a single sample with missingness in excess of 20%. All remaining samples exhibited high consistency between tightly linked markers: log odds ratio error scores were less than 2.0 for all samples (Lincoln and Lander (1992)). The final set of genetic data consisted of 946 mice.

For each DO mouse, we compared its genotype to that of the eight founder strains at all 110,807 markers to calculate the probability that a given founder contributed a given allele at that marker (implemented in the R package: qtl2 Broman *et al*. (2019)). In other words, the founder-of-origin probability is the likelihood a given DO mouse possess a specific founder haplotype at the focal marker and can be used to identify genomic regions that are identical-by-decent. This allowed us to directly test for an association between the founder-of-origin probability and phenotype at all genotyped markers. Using the founder-of-origin of consecutive typed markers and the genotypes of untyped variants (SNPs and small insertion-deletions) in the founder strains, we then imputed the genotypes of all untyped variants (34.5 million) in all 946 mice. The majority, but not all, of imputed variants were bi-allelic SNPs. Targeted association testing at imputed variants allowed us to fine-map many QTLs to near single gene resolution.

### Genetic Linkage Analysis

With the R qtl2 package, we calculated kinship matrices using the leave-one-chromosome-out (LOCO) method and conducted quantitative trait locus mapping analyses (Broman *et al*. (2019)). In order to identify significant additive genetic associations, we fit a linear mixed model with diet and founder-of-origin probabilities per marker as fixed effects and kinship as a random effect. To identify significant genotype by diet (GxD) interaction effects, we fit a linear mixed model with diet and founder-of-origin probabilities and their interactions as fixed effects and kinship as a random effect. To calculate an LOD score for the GxD interaction term we subtracted the LOD score of the full model from the additive model. To determine whether the interaction LOD score was statistically significant, we conducted a permutation analysis by randomizing phenotype values (regardless of dietary treatment), fitting both the full and the additive models, subtracting the genome wide set of LOD scores of the full model from the additive model and storing the maximum LOD value (Churchill and Doerge (1994)). We repeated this procedure 1,000 times to obtain a distribution of maximum LOD scores and applied empirical p-value threshold of 0.05 to define significant QTLs and 0.1 as suggestive QTLs.

For each significant and suggestive QTL, we imputed variants for 5Mb +/-the lead marker position and re-ran the QTL mapping procedure (implemented in the snpscan function from qtl2). To assess the significance of imputed variants for each region, we re-ran the permutation procedures as previously described with 1,000 iterations and applied empirical p-value threshold of 0.05. Finally, we identified all candidate variants as those that are specific to lead founder-allele-pattern (FAP), or if the lead FAP contains fewer than 10 variants, we also include variants specific to the second ranked FAP. We identified lead candidate genes by their proximity to candidate FAP variants and by cross-referencing against phenotypic effect in the Mouse Genome Informatics (www.informatics.jax.org) database.

### Heritability and Genetic Correlations Analyses

For each trait, we calculated the additive genetic variance relative to phenotypic variance, e.g. narrow-sense heritability, and its 95% credible interval using a Bayesian model with diet as a fixed effect and kinship as a random effect based on the EMMA model as implemented in R’s STAN package (Kang *et al*. (2008); Carpenter *et al*. (2017); Stan Development Team (2020)). We assessed whether heritability was significantly greater than zero by applying one-sided z-test to the posterior distribution with false discovery rate controlled at 0.05.

To measure the degree to which the additive genetic variance underlying two traits is shared we calculated their genetic correlation using the mathematical framework described in Furlotte and Eskin (2015). We used a Bayesian model implemented in R’s STAN package (Stan Development Team (2020)) to estimate the genetic correlation and its 95% credible interval. The details about model assumptions and priors are in the Supplemental Materials. We ran three independent chains with 2,000 Markov chain Monte Carlo (MCMC) iterations, and posterior estimates were derived by combining all three MCMC chains after 1,000 burn-ins. The convergence diagnostics were assessed by Gelman-Rubin’s statistic (Gelman and Rubin (1992)). The significance of phenotypic correlation was determined by t-test and the significance of genetic correlation was determined by posterior mean and standard deviation under standard normal distribution. We applied Benjamini and Hochberg (1995) method to control significant phenotypic and genetic correlations respectively, at a false discovery rate of 0.05.

### Comparison of full and reduced genotype-by-diet association models to measure interaction effects

In order to determine which diet intervention(s) are responsible for genotype-by-diet interaction effects, we re-tested the lead genotyped marker at each significant QTL in the following models:

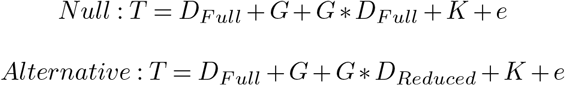

where *T* is trait, *G* is genotype, *K* is kinship, *e* is error, *D*_*Full*_ is all five treatments and *D*_*Reduced*_ eliminates, in singles or pairs, 1D fast, 2D fast, 20% CR, or 40% CR. We first remove a single diet at a time and evaluate the fit of each alternative model using the likelihood ratio test. The diet with the highest LOD score is then tested in pairs with each of the other three diets to determine whether model fit is improved.

## Results

### Dietary intervention altered physiology of early adulthood mice

We measured the effect of dietary interventions on multiple aspects of mouse physiology and found that both the type (CR vs IF) and magnitude of each intervention affected the physiological response. To summarize, the 40% CR intervention had the greatest impact, 24 of 36 total traits were significantly different compared to the AL diet (Figure 2). For select traits we also provide the non z-score transformed values (Supplemental Figure S1). Following the 40% CR intervention, the 20% CR, 2D fast, and 1D fast treatments resulted in 11, 9 and 4, traits changing significantly in comparison to the AL group (Figure 2). Examining body weight, body length, percent lean mass, tissue mass, tissue area, and bone mineral content, the treatment with the largest effect in comparison to AL was 40% CR and this effect was more than double the difference between 20% CR and AL (Figure 2). Interestingly, the 2D fast and 20% CR had nearly the same mean body weights, however the treatments exhibited opposite effects on body fat percentage: 2D fast reduced and 20% CR increased DX_PFAT(Figure 2). In summary, we found intermittent fasting and daily caloric restriction had distinct effects on multiple body and bone composition traits and changes in response to the 40% CR and 2D fast treatments were not simply a doubling of magnitude of 20% CR and 1D fast treatment effects. These patterns were also observed for additional physiological traits.

**Fig. 2.**
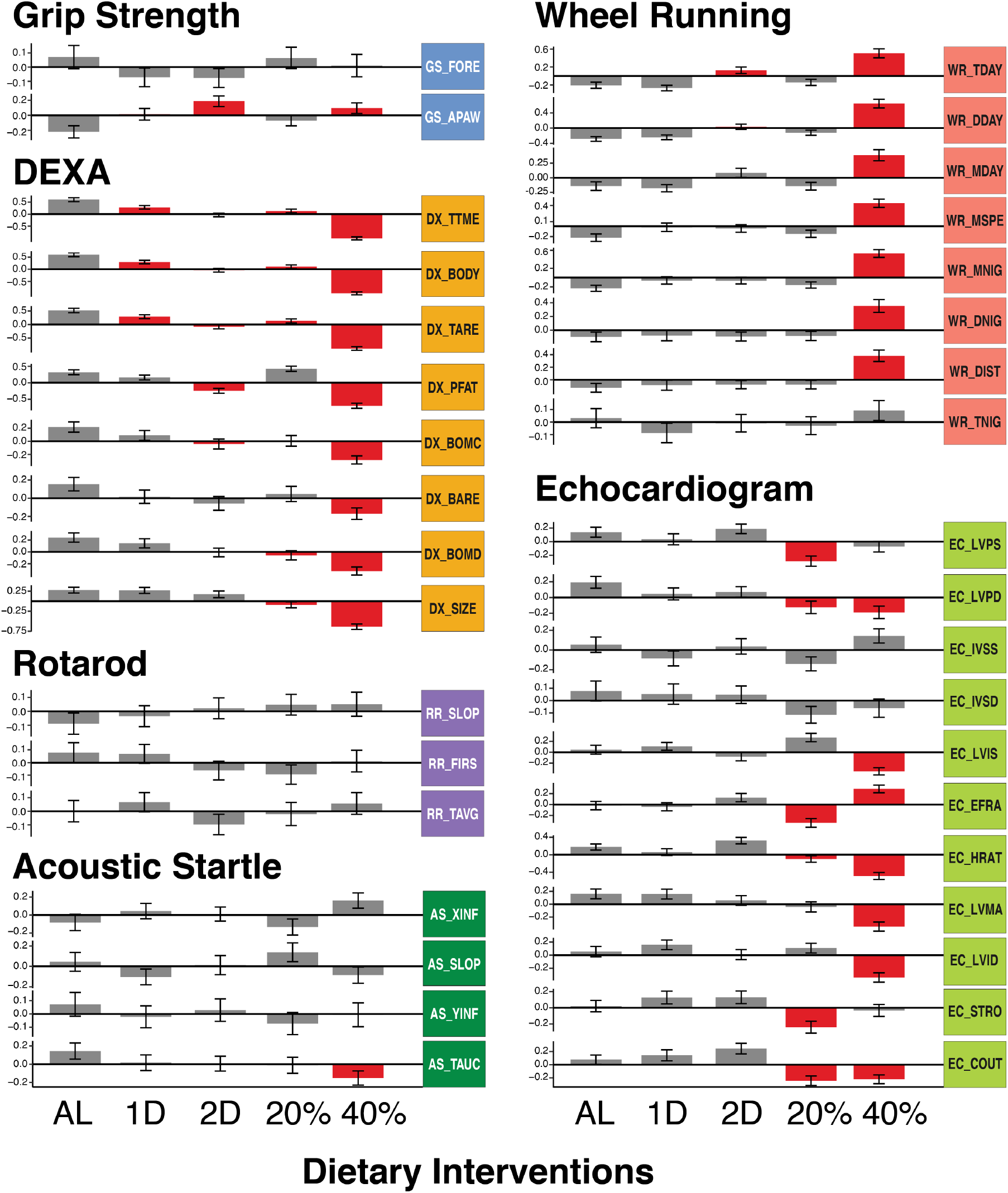
Diet specific mean (SE) trait values for all experimental procedures. All trait values were z-score transformed following batch and generation correction. Red bars denote traits that were significantly different from AL diet.

We uncovered multiple cardiac phenotypes which were significantly altered by both the 20% CR and 40% CR treatments whereas no significant effect was detected in the intermittent fasting treatments. Heart rate, cardiac output, and diastolic left ventricle wall thickness (EC_HRAT, EC_COUT, EC_LVPD) were significantly lower in both CR groups compared to AL (Figure 2). Additionally, the 20% CR group exhibited significantly lower systolic left ventricle wall thickness (EC_LVPS) and stroke volume (EC_STRO) whereas the 40% CR group exhibited significantly lower left ventricle mass and inner dimension in systole and diastole (EC_LVMA, EC_LVIS, EC_LVID). The cumulative effect of these divergent responses was that the 20% CR group had the lowest ejection fraction and the 40% CR group had the highest ejection fraction (Figure 2, EC_EFRA). Similarly, after controlling for body weight, we found cardiac output was lowest for the 20% CR group and highest for the 40% CR group (Supplemental Figure S1E,F). These results suggest that caloric restriction, and not intermittent fasting, was detrimental to the cardiovascular efficiency of early adulthood mice treated with 20% CR and beneficial to the 40% CR group. This pattern of effects was similar to the effects on lean and fat mass, in which 20% CR, and 40% CR treatment effects relative to AL varied in both magnitude and sign (positive or negative).

We conducted multiple experiments to measure neuromuscular and motor function: running on a wheel, grip strength, and balancing on the rotarod. Wheel running activity, measured as total distance, max speed, and amount of time on a running wheel, were significantly increased in the 40% CR treatment compared to all other groups for both the light and dark cycles (Figure 2). The 2D fast treatment exhibited a significant increase in total wheel time and moderate increases in distance and max speed during the day compared to the other groups (Figure 2). No wheel running traits were significantly different in the 1D fast or 20% CR treatments in comparison to the AL group (Figure 2). The only significant difference observed among the grip strength and rotarod traits was an increase in all-paws grip strength in the 40% CR and 2D fast treatments (Figure 2). To summarize the effect of dietary intervention on neuromuscular and motor function, the 40% CR treatment, followed by 2D fast, ran the farthest, and -by at least one measure-had the greatest strength. Interestingly, these same groups had the lowest body weight, lowest body fat percentage, and highest lean mass percentage.

We measured hearing ability using the acoustic startle experiment. We found the AL treatment had the most sensitive hearing whereas the 40% CR treatment mice had the least sensitive hearing, when measured as the total area under the startle response curve (AS_TAUC, Figure 2). This result suggested that 6 month exposure to 40% CR treatment, in contrast to all other interventions, had a detrimental effect on hearing ability in 12 month old mice.

Collectively, these results demonstrated that intermittent fasting and daily caloric restriction had distinct effects on multiple aspects of physiology and neither the magnitude nor the sign of effects were linear with respect to daily calorie intake or length of intermittent fasting regime. Additionally, none of these interventions were universally beneficial across all aspects of organismal physiology. Finally, we found the effect of dietary intervention was correlated between many traits. In some instances, this was because one trait was directly calculated from another trait measured in the same assay (see Methods: Phenotypic Assays). Alternative and mutually non-exclusive hypotheses may also explain these results: 1) the traits measured similar aspects of physiology (e.g. fat mass and body weight), 2) the traits were derived from the same phenotypic assay and environmental variables (e.g. time of day, time of year, experimenter) were constant, and 3) trait variation is controlled by a shared genetic basis. In order to investigate these hypotheses, we estimated the heritability of each trait and their pairwise phenotypic and genetic correlations.

### The majority of physiological traits exhibit significant genetic heritability

To determine the contribution of genetics to phenotypic variation in each trait irrespective of diet, we calculated heritability across all animals in the study and found that most traits measured at one year of age (31 of 36) have significant heritability (Figure 3). Body composition traits from DEXA and one measure of hearing sensitivity had the highest heritabilities (>0.5). Wheel running speed and distance traits had moderate (0.3-0.5) heritabilities. Several cardiac traits, including heart rate, stroke volume and cardiac output, as well as forepaw grip strength and time to fall on the rotarod had low (0.1-0.3) but statistically significant heritabilities. Traits with heritabilty not significantly different from zero included two echocardiogram derived traits, and one each for acoustic startle, rotarod, and grip strength (Figure 3).

**Fig. 3.**
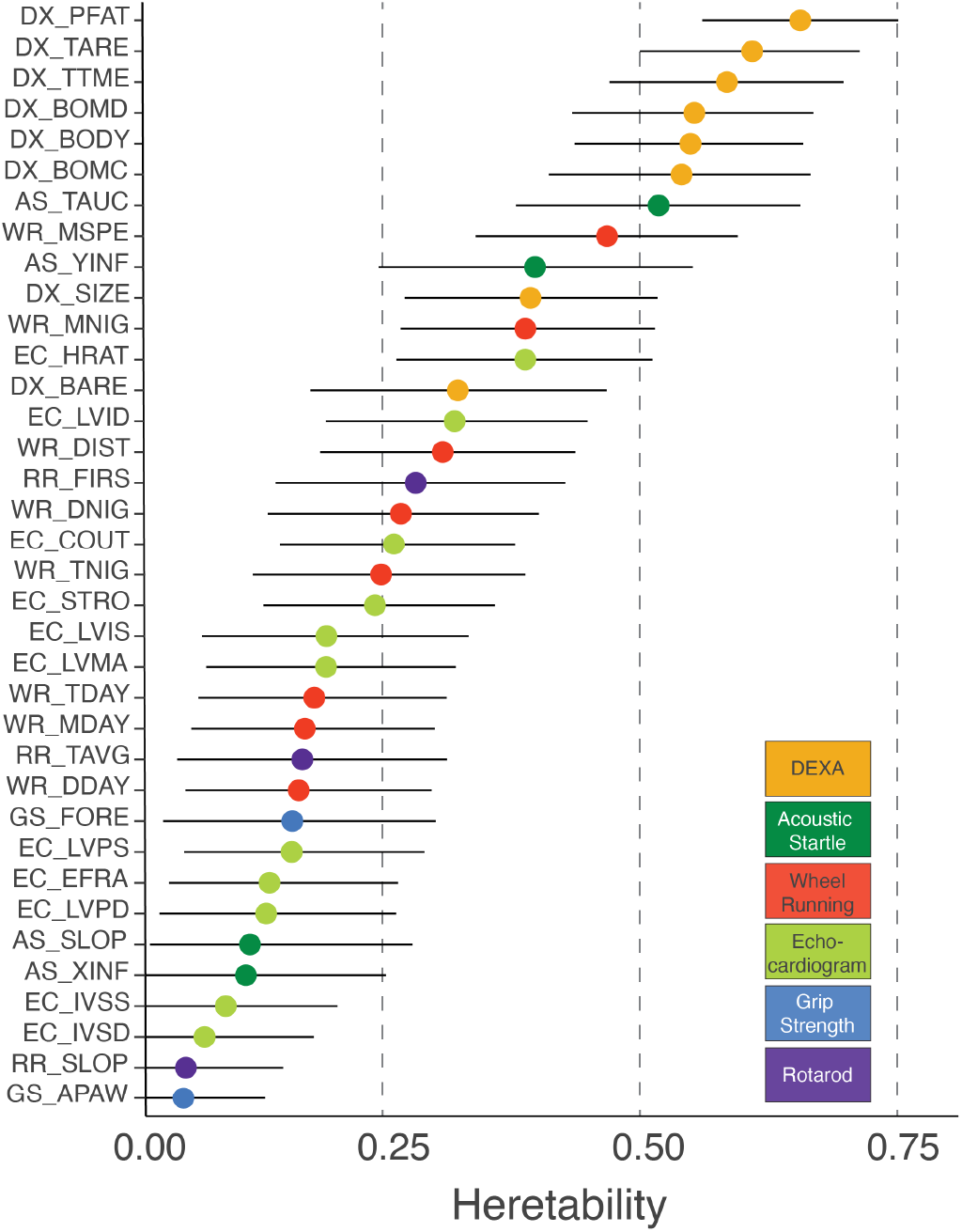
Trait specific heritability (95% Bayesian credible interval) values.

### Phenotypic and genetic correlations separate distinct aspects of physiology

We calculated the phenotypic correlation between all trait pairs using all samples and found that many trait pairs, especially those derived from the same assay, were tightly correlated. (Figure 4A, lower-triangle). We also calculated diet-specific correlations and found these to be very similar to correlations obtained when using all animals (Supplemental Figure S2). When diet-specific differences were observed, they affected the magnitude but not the sign of the correlation. For example, cardiac output and stroke volume (EC_COUT, EC_STRO) were positively correlated with body composition traits (DX_PFAT, DX_TARE, DX_BODY, and DX_TTME) in AL, 1D and 2D group, however, the correlation was reduced in 20% CR and 40% CR groups (Supplemental Figure S2). Since the phenotypic correlations were largely similar across diets, we used correlations calculated from all animals in subsequent analyses.

**Fig. 4.**
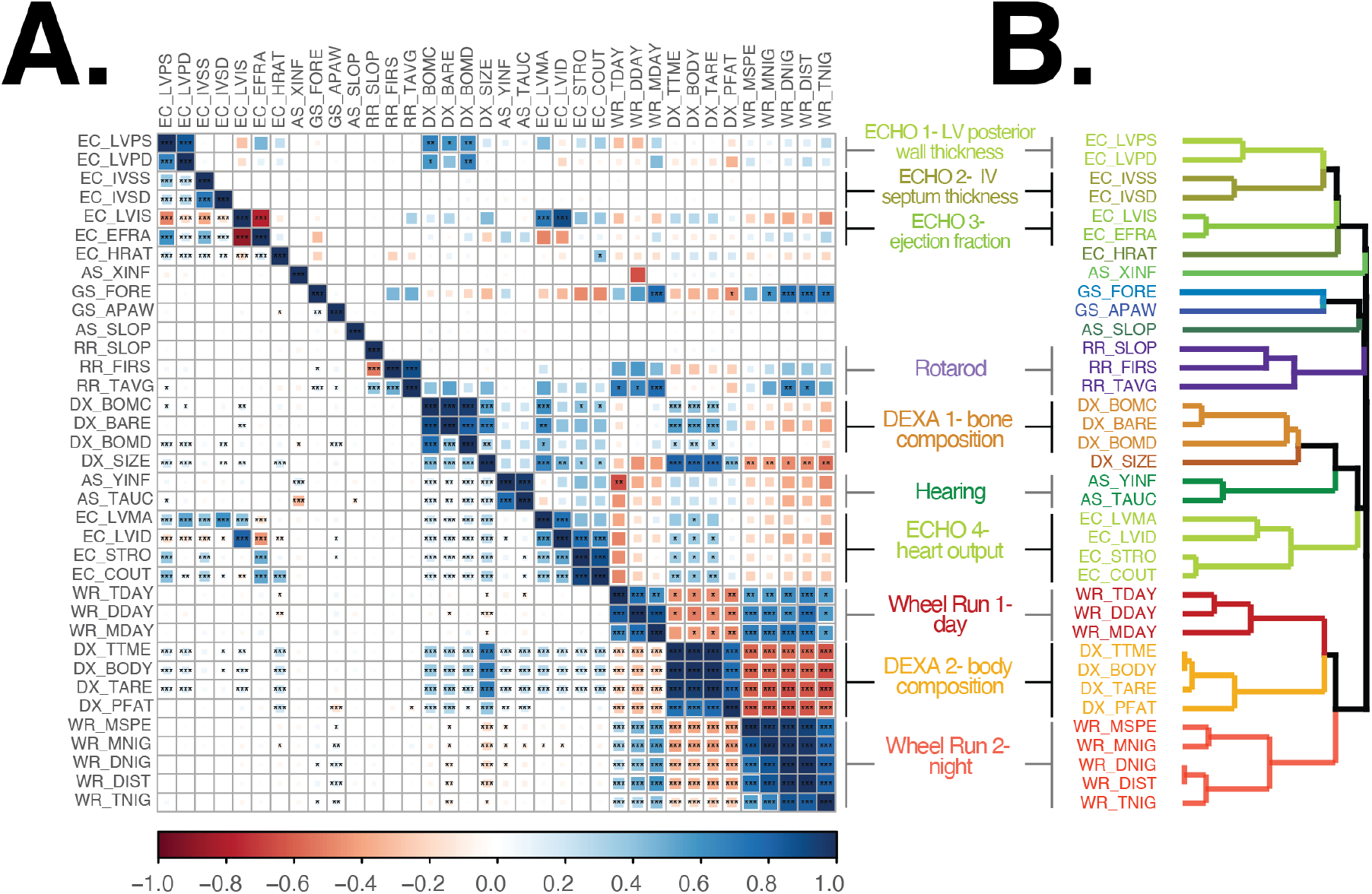
A. Pairwise genetic (upper-triangle) and phenotypic (lower-triangle) correlations. * p-value <0.05. ** p-value < 0.01, *** p-value < 0.001 (FDR adjusted p-value) B. Hierarchical clustering of traits used phenotypic correlation values. Each color represents a significantly distinct cluster.

In order to measure the degree to which the heritable fraction of variation in two traits was shared we calculated their genetic correlation (Figure 4A, upper-triangle). This value measures the correlation of genetic effects on two traits, and a genetic correlation equal to one means that every variant that affects the first trait has an equal pleiotropic effect on the second trait. For many traits, the genetic and phenotypic correlations were similar (adjusted R square of 0.62, Supplemental Figure S3). Additionally, we identified 138 instances (out of 630 trait pairs) for which the phenotypic correlation was significantly greater than zero but the estimated genetic correlation was indistinguishable from zero. This suggested that, for these trait pairs, the phenotypic correlation was due to shared environmental factors.

We sought to quantify the degree of similarity between traits using an unsupervised hierarchical clustering analysis of all pairwise phenotypic correlations. We identified 10 clusters of two or more traits and six single-trait clusters (Figure 4B). All 10 multi-trait clusters were composed of traits from the same assay, however traits from all assays (except rotarod) were split across multiple clusters in non-adjacent regions of the dendrogram (Figure 4B). For example, DEXA derived body composition traits formed two multi-trait clusters, the first cluster was composed of bone physiology traits and was adjacent to a cardiac output cluster, whereas the second cluster was composed of body area/tissue composition traits and was located within day and night time wheel running clusters (Figure 4B). We interpret traits in distinct clusters as measurements of distinct aspects of physiology, with cluster placement in the dendrogram indicating the degree of similarity between these aspects of physiology.

Multiple factors may contribute to the high correlations within each multi-trait cluster: different traits measured the same underlying physiology, the shared environment in which traits were measured, and a shared genetic basis. In eight of 10 multi-trait clusters, nearly all trait pairs within each cluster were significantly genetically correlated with each other (Figure 4A), suggesting that the traits that comprise these aspects of physiology shared a common genetic basis. In the two remaining multi-trait clusters (Rotarod and ECHO 2), trait pairs were, for the most part, not significantly genetically correlated (Figure 4A) because of the low genetic heritability of one or both traits (Figure 3). This result suggested that the significant phenotypic correlations within these clusters was likely due to shared environmental factors. We next sought to measure the diet-independent and diet-dependent genetic basis of each directly measured trait using a QTL mapping approach.

### Genetic mapping with founder-allele-patterns identifies candidate variants

Using both additive and genotype-by-diet (GxD) interaction models, we used the founder-of-origin genotype probabilities to map associations for each of the 36 directly measured traits. For the additive model, we found 16 significant QTLs (p-value < 0.05) and seven suggestive QTLs (p-value < 0.1) among the 36 phenotypic traits (Table 1). In instances in which multiple traits map to the same genomic region we count these as a single QTL. For the GxD interaction model, we identified two significant QTLs-both of which were associated with cardiac physiology traits- and one suggestive QTL for hearing sensitivity (Table 1).

**Table 1.**
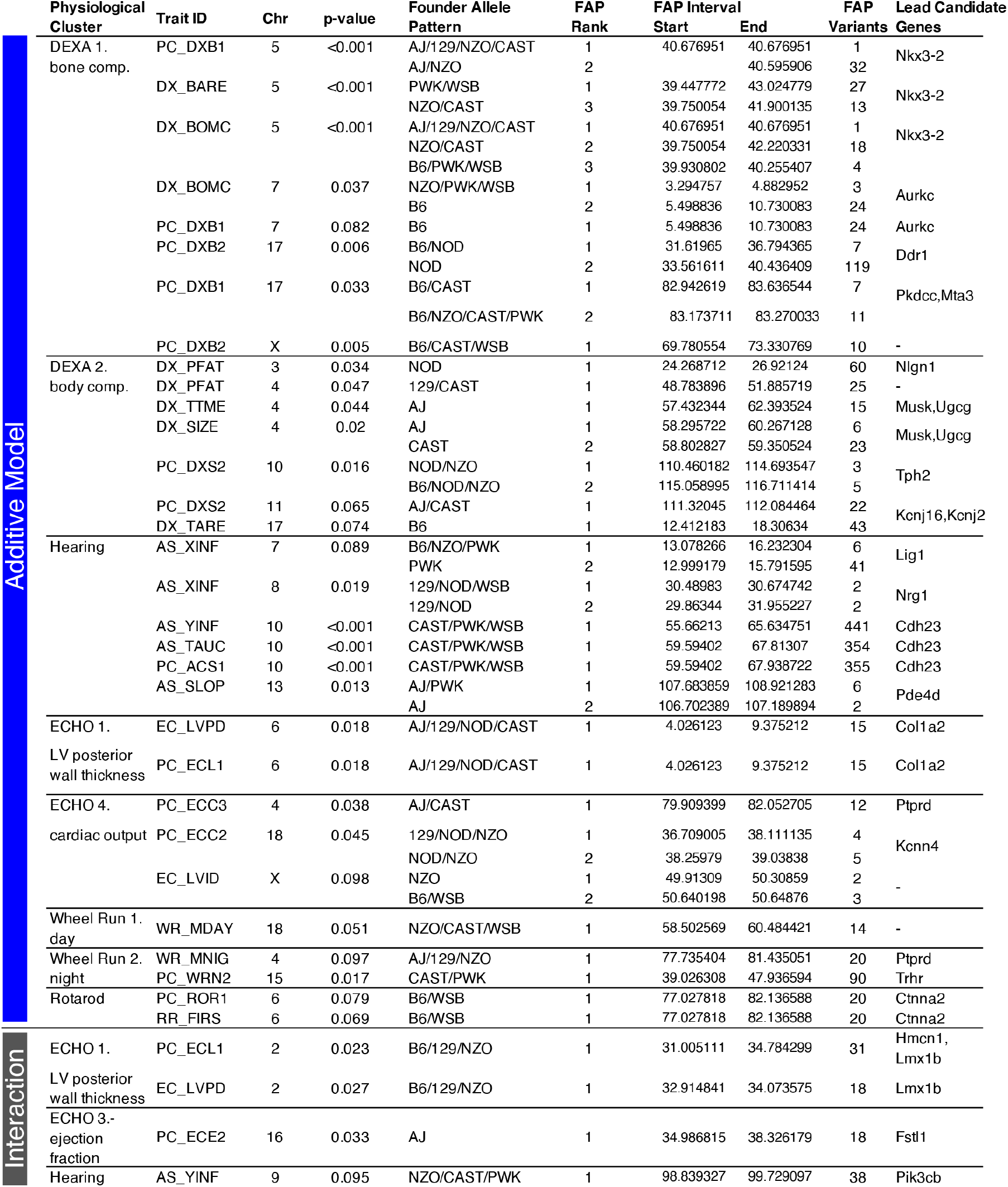
Genome-wide significant diet-independent and diet-dependent QTLs. Traits are organized by clusters identified in Figure 4. For each trait, we calculated a genome-wide significant LOD score threshold using a permutation analysis. We identified the FAP of the variant with the strongest LOD score, the genomic location of these variants, and the number of significant variants that comprise the lead FAP group. For loci in which the lead FAP is comprised of fewer than 10 variants, we also present results for the second ranked FAP. We list likely candidate genes based on lead FAP variants and a survey of gene knock-out phenotypes.

To more thoroughly interrogate aspects of physiology represented by each multi-trait cluster, we conducted a principal component analysis of the traits in each cluster (Supplemental Table S3) and repeated the genetic association analyses. For the PC derived traits, we identified eight diet-independent and one diet-dependent QTLs that were not identified in our analysis of the directly measured traits (Table 1).

To more precisely fine-map the genomic interval of each QTL, we imputed all SNPs and small insertion-deletion variants from the fully sequenced DO founders (Keane *et al*. (2011)) across a 5Mb interval centered at the lead genotyped marker and used these variants to conduct the fine-mapping association analysis. For each imputed variant, we identified the founder-of-origin for the major and minor allele Wright *et al*. (2020). To illustrate this process, consider a bi-allelic A/G variant, if allele A was specific to founders AJ, NZO, and PWK and allele G was specific to the other 5 founders, then we assigned A to be the minor allele and defined a founder-allele-pattern (FAP) of AJ/NZO/PWK for this variant. Importantly, the FAP is a measure of identity-by-state for imputed SNPs, and contrasts with the founder-of-origin genotype probabilities that measure identity-by-decent in the DO population.

To identify the variants and founder haplotypes most likely responsible for the association at each locus, we grouped variants based on their FAP and ranked groups based on the largest LOD score among its constituent variants. (Note that, by definition, no variant can be a member of more than one FAP group.) We hypothesized that the functional variant(s) responsible for trait-specific variation were among those in the lead FAP group because they exhibit the strongest statistical association and it is unlikely any additional variants are segregating in this genomic interval beyond those identified in the full genome sequences of the eight founder strains. By focusing on FAP groups with the largest LOD scores, we narrowed the number of candidate variants at each QTL. The lead FAP and the number and location of statistically significant variants that comprise each FAP group are summarized in Table 1. Additionally, a list of all imputed variants significantly associated with each trait and the candidate genes in each region are provided in Supplemental Files 1 and 2. To demonstrate this approach, we fine-mapped QTLs associated with bone composition traits.

### Alleles of contrasting effects associated with variation in bone composition

Traits comprising the tightly correlated bone composition cluster (Figure 4), were associated with a chromosome 5 locus (total bone area and mineral content) and a chromosome 7 locus (bone mineral content, Figure 5A). It is unsurprising that the locus with the greatest LOD score, chromosome 5, was associated with both total bone area and bone mineral content because these two traits are both correlated with mouse size (Brommage (2003)). We repeated the genetic association analysis with PC derived bone composition traits and found PC1 replicated the chromosome 5 association and the strength of the chromosome 7 association was reduced (Table 1). Additionally, the PC1 analysis identified a new peak on chromosome 17 and PC2 analysis identified two new QTLs on chromosomes 17 and X (Figure 5B). To identify candidate variants and genes, we fine-mapped these loci using the FAP group of each imputed variant.

**Fig. 5.**
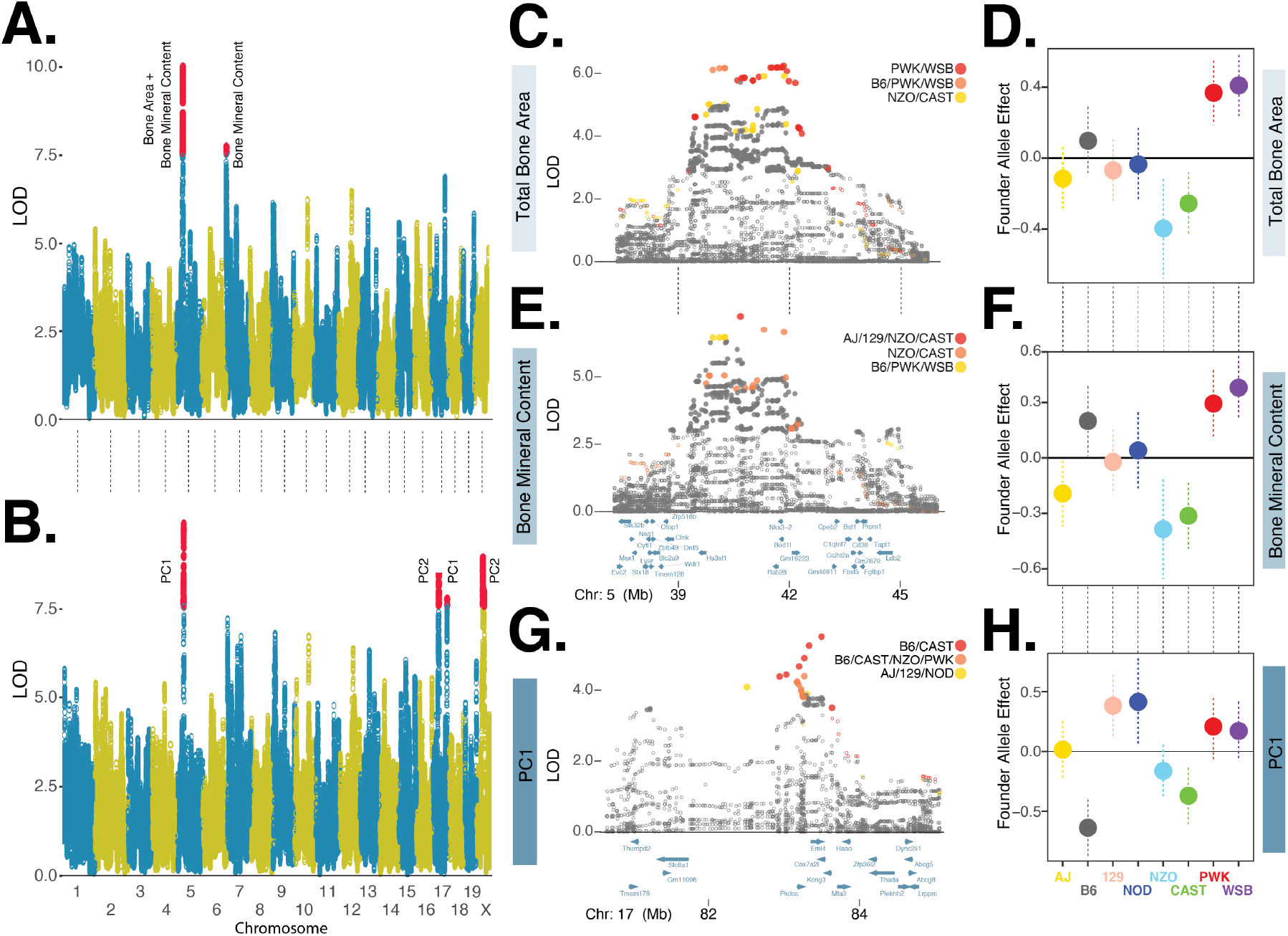
A. Manhattan plot of directly measured bone composition traits: total bone area and bone mineral content. Red circles denote markers with statistically significant (p < 0.05) LOD score based on genome-wide permutation analysis. B. Manhattan plot of PC derived bone physiology traits. C. Fine mapping of total bone area chromosome 5 locus using imputed variants. LOD scores of closed circles are statistically significant (p < 0.05) based on permutation analysis of all imputed variants with +/-5Mb of lead genotyped marker. Variants in three founder-allele-pattern groups shown in red, orange, and yellow circles, ordered by maximum LOD score. D. Founder allele effects and standard error estimates for the lead genotyped variant for total bone area. E. and F. are for bone mineral content, details are the same as C. and D. G. and H are for the bone composition-PC1 chromosome 17 locus, details are the same as in C. and D.

We fine-mapped the chromosome 5 loci associated with total bone area (DX_BARE) and bone mineral content (DX_BOMC). The two lead FAPs-ranked by maximum LOD score of each FAP variant group-for DX_BARE contained variants with minor alleles specific to the PWK and WSB founders and the rank 3 FAP was comprised of NZO and CAST (Figure 5C). The PWK and WSB alleles were associated with the largest positive effect of the eight founders on total bone area, whereas NZO and CAST were associated with the largest negative effect (Figure 5C). Next, we examined the fine-mapping results for DX_BOMC and identified a different order of lead FAPs: rank 1 and 2 groups contained minor alleles specific to the NZO and CAST founders whereas the rank 3 group was comprised of PWK and WSB (Figure 5D). The effect of the founder alleles on bone mineral content (Figure 5E) were similar to results for total bone area (Figure 5C). Although the rank order of the top three FAP groups differed slightly between the two traits, these results are consistent with the hypothesis that this one locus affects these two highly similar traits. Moreover, we have identified at least three distinct alleles at this locus: a positive allele derived from the PWK and WSB founders, a negative allele derived from the NZO and CAST founders, and a neutral allele derived from the four other founders.

To further illustrate the utility of fine mapping QTLs with variants grouped by FAP, we examined the chromosome 17 locus associated with bone composition PC1 (Figure 5B). We found that variants with minor alleles specific to the B6 and CAST founders exhibited the strongest statistical association (Figure 5G). Consistent with the composition of this lead FAP, we found the effect of the B6 and CAST founder alleles to have the largest negative effects on bone composition PC1 (Figure 5H). We next used FAP grouped variants and the Mouse Genome Informatics database of phenotypic effects (www.informatics.jax.org) to identify candidate genes.

The chromosome 5 total bone area QTL contained 406 significant variants, of which 27 (21 intergenic SNPs, 6 intronic SNPs) were specific to the positive effect PWK/WSB alleles (rank 1 FAP group) and 13 (12 intergenic SNPs and 1 intronic SNP) were specific to the negative effect NZO/CAST alleles (rank 3 FAP group; Table 1, Figure 5C). The chromosome 5 bone mineral content QTL contained 350 significant variants, of which 4 intergenic SNPs were specific to the positive B6/PWK/WSB alleles (rank 3 FAP group) and 19 (17 intergenic SNPs and 2 intronic SNPs) were specific to the negative effect NZO/CAST alleles (rank 1 and 2 FAP groups; Figure 5E). We found no protein coding variants in the lead FAP groups that were significantly associated with either trait, suggesting that the functional variant(s) altered gene expression. Many candidate variants were located in intergenic regions adjacent to *Nkx3-2* (Figure 5C,E), which encodes a homeobox protein critical to skeleton development (Lettice *et al*. (1999)). The chromosome 17 locus associated with bone composition PC1, was comprised of 47 statistically significant variants, seven of which were members of the B6/CAST FAP (Figure 5G). All of these variants were intergenic SNPs located in a genomic interval containing five genes (*Pkdcc, Eml4, Cox7a21, Kcng3*, and *Mta3*) and of these candidates *Pkdcc* has previously been shown to effect bone morphology (Sajan *et al*. (2019); Imuta *et al*. (2009)).

These analyses illustrate three key findings: 1) conducting genetic association analyses with both directly measured and PC derived traits can reveal novel loci, 2) fine mapping loci with FAP groups greatly reduces the number of lead candidate variants, and 3) FAP variant groups illuminate the link between specific founder haplotypes associated with positive, neutral, or negative phenotypic effects.

### Cardiac physiology is altered in response to dietary intervention in a genotype dependent manner

All three significant diet-dependent QTLs were associated with cardiac physiology traits (Table 1). We identified one QTL associated with the second PC (PC2_ECE2) of ejection fraction (EC_EFRA) and left ventricular inner dimension, systole (EC_LVIS) (Figure 6A). These two traits are positively correlated with PC2_ECE2 (Supplemental Figure S4), which we interpreted as a measure of heart pumping efficiency. We fine-mapped this QTL and found the lead FAP was composed of AJ-specific alleles (Figure 6A). The remaining QTLs were associated with diastolic left ventricular posterior wall thickness (EC_LVPD) and the first principal component (PC_ECL1) of EC_LVPD and EC_LVPS, systolic left ventricular posterior wall thickness (Table 1). EC_LVPD and EC_LVPS are positively correlated (Figure 4A) and, unsurprisingly, the QTLs for PC_ECL1 and EC_LVPD were located in the same region of chromosome 2 and shared the same lead FAP: B6/129/NZO (Figure 6B,C). We found the genomic interval associated with PC_ECL1 to be larger than EC_LVPD (30.9-34.8Mb versus 32.9-34.1Mb) and fine-mapping EC_LVPS revealed a region of association between 30.5 and 32.0 Mb (Supplemental Figure S5). Although the size of our mapping population limits our ability to conclude whether the associations with systolic and diastolic wall thickness are separate loci affected by distinct functional variants, this result does explain the subtle difference between the fine mapped intervals for PC_ECL1 and EC_LVPD (Figure 6B,C). We next set out to determine which dietary intervention(s) were responsible for these genotype-by-diet interaction effects.

**Fig. 6.**
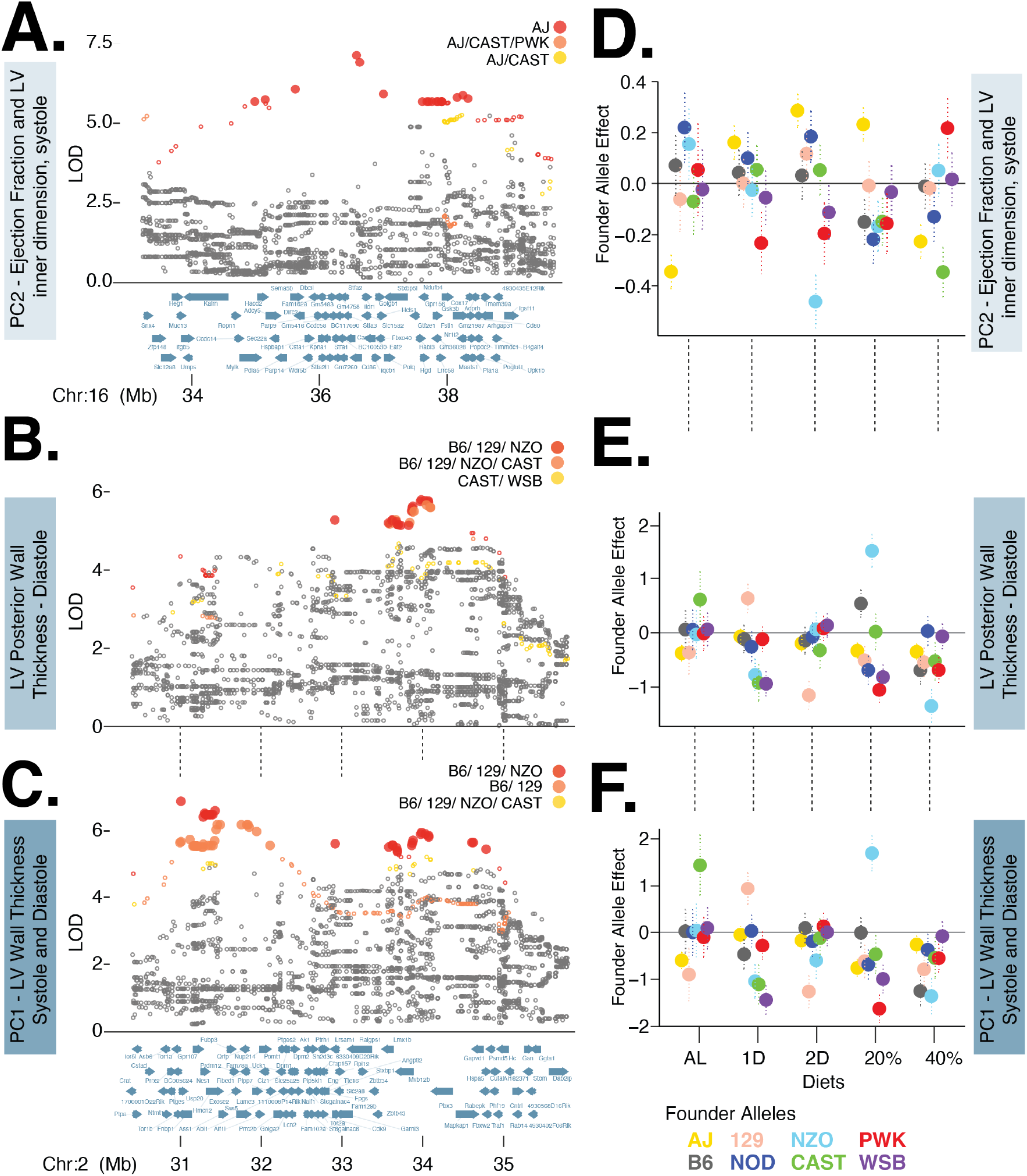
A. Fine mapping of chromosome 16 locus associated with PC2 of ejection fraction and left ventricular inner dimension, systole. Rank 1, 2, and 3 FAP variants shown in red, orange, and yellow circles. LOD scores of closed circles are statistically significant (p < 0.05) based on permutation analysis of all imputed variants with +/-5Mb of lead genotyped marker. B. Fine mapping of chromosome 2 locus associated with left ventricular posterior wall thickness, systole. Legend same as A. C. Fine mapping of chromosome 2 locus associated with PC1 of left ventricular posterior wall thickness, systole and diastole. Legend same as A. D-F. Diet-specific effect of lead genotyped variant for each of the eight founder variants for three focal traits.

In order to determine the diet most likely responsible for the significant GxD interaction effects, we compared the lead variant LOD score in the full model to reduced models in which we pruned diets in singles and pairs. We considered a diet as likely responsible for the significant interaction effect if the removal of that diet reduced the strength of the association in comparison to the full model. For PC_ECE2, we found that 20% CR and 2D fast treatments were most likely responsible for the diet-dependent association (Supplemental Table S4). The diet-specific founder-allele effect for the AJ allele exhibited the largest positive effects in the 20% CR and 2D fast treatments and significant negative effects in AL and 40% CR treatments (Figure 6D). These results are consistent with the hypothesis that the diet-dependent effects of the AJ allele were responsible for the interaction association at this locus.

We identified 18 variants significantly associated with PC_ECE2 and all of these were specific to the lead FAP, AJ. A single variant was an intergenic structural variant, and the remaining 17 were non-coding exonic (1), intronic (4) or intergenic (12) located at nine genes. One variant was located in the 3’ UTR of Follistatin-like 1 (*Fstl1*), this is a secreted glycoprotein expressed in the adult heart that affects cardiac morphology, contractility, and vascularization (Oshima *et al*. (2008); Shimano *et al*. (2011)).

We next examined the diet-dependent associations with EC_LVPD and PC_ECL1 and, using the reduced GxD association model test, found that 20% CR and 1D fast treatments were most likely responsible for this interaction QTL (Supplemental Table S4). We estimated the diet-specific founder-allele effects for the lead variant at this QTL and focused on the effects of B6, 129 and NZO. We estimated distinct diet-specific effects for each founder: the effect of B6 was significantly negative in 40% CR, positive in 20% CR (for EC_LVPD only), and largely neutral in the other three diets; the effect of 129 was significantly positive in the 1D fast treatment and negative in the other four diets; the effect of NZO was significantly positive in 20% CR, neutral in AL, and negative in the other three diets (Figure 6E,F). Although the B6 allele was identified in the lead FAP, the effect size results suggest this allele was unlikely to be responsible for the GxD interaction association. The seeming incongruence between the FAP and effect-size estimates illustrates a key point: FAPs were annotated using imputed variants and reflect identity-by-state whereas effect-sizes were estimated using the founder-of-origin probabilities and reflect identity-by-descent (as described in Methods section). These results were consistent with the hypothesis that either 129 or NZO founder alleles were responsible for the significant interaction QTL because of the strong diet-specific effect of the 129 allele in 1D fast and NZO allele in 20% CR. Additionally, these results would be consistent with the hypothesis that both founder alleles are responsible and, given our observation of contrasting diet specific effects, each may harbor distinct functional variants at this QTL.

We identified a total of 59 and 28 variants significantly associated with PC_ECL1 and EC_LVPD. Thirty one PC_ECL1 variants and 18 EC_LVPD variants were specific to the lead FAP (B6/129/NZO) and all variants were SNPs. Variants associated with PC_ECL1 (19 intronic and 12 intergenic) were located in close proximity to 10 genes. Ten variants (eight intronic and two immediately upstream) were located at Hemi-centin2, a fibulin family extracellular matrix protein. Genetic knock-out studies of *Hmcn2* have resulted in abnormal left ventrical morpholgy in mice (Dickinson *et al*. (2016)) and have been associated with electrocardiogram derived traits in humans (Tereshchenko *et al*. (????)). Additionally, three variants (intronic) were located at *Lmx1b*, a LIM homeobox transcription factor 1-beta that is known to regulate limb and organ development (Dreyer *et al*. (1998); Schweizer *et al*. (2004); Doucet-Beaupré *et al*. (2016)). Variants associated with EC_LVPD (9 intronic and 9 intergenic) were located in close proximity to 5 genes, a list which included *Lmx1b* and lacked *Hmcn2*.

Taken together, all significant diet-dependent QTLs were associated with heart physiology. Fine mapping with FAPs narrowed the likely number of causal variants and identified candidate genes previously linked to cardiac morphology or function. We previously showed that the signs of the effects of diet on the mean physiological trait measurements were specific to the type of intervention (CR or IF) and their magnitudes were non-additive with respect to the magnitude of intervention (Figure 2). Identification of candidate genes with diet-dependent effects suggests molecular mechanisms to explain these results.

## Discussion

### Conditionally beneficial effects of CR and IF on distinct aspects of physiology

A primary goal of this study was to address the question: which aspects of physiology would respond to dietary intervention in early adulthood mice? We performed this experiment using DO mice in order to assess this question in an outbred genetic model that more closely resembles human populations. Additionally, we were interested in determining whether the physiological health benefits (or detriments) of daily CR could be replicated with intermittent fasting treatments. We found dietary intervention initiated at six months of age significantly altered many traits in 12 month old mice. The 40% CR dietary intervention impacted the greatest number of traits in comparison to the AL diet followed by the 20% CR, 2D and 1D fast treatments (Figure 2). Using six experimental assays we clustered individual traits into distinct aspects of physiology (Figure 4). In many instances, changes to an aspect of physiology were not consistent between IF and CR. For the body composition cluster, we observed similar mean body weights for the 20% CR and 2D fast treatments, however the 2D fast treatment significantly increased the proportion of lean muscle mass and reduced fat mass, whereas the 20% CR decreased the percentage of lean muscle mass and increased fat mass (Figure 2). How might these changes in physiology impact organismal health?

While the lifespan extension of daily CR is well established, it remains largely unknown whether dietary intervention would improve physiological function in healthy, early adulthood mice. Our results demonstrated that 2D fast and 40% CR, in comparison to 20% CR, improved multiple aspects of cardiovascular function. Left ventricular posterior wall thickness (systolic and diastolic) increased in 2D fast but decreased in 20% CR, and these changes in morphology were correlated with cardiac function - ejection fraction and stroke volume increased in 2D fast and decreased in 20% CR (Figure 2). Similar to the 2D fast treatment, we observed a decrease in posterior wall thickness for 40% CR and an increase in cardiac function - measured as increased ejection fraction and stroke volume after controlling for the dramatic decrease in body weight observed in the 40% CR mice (Figure 2, Supplemental Figure S1F). Ejection fraction was previously shown to decrease with mouse age and is indicative of decreased cardiac health (Medrano *et al*. (2016); Lindsey *et al*. (2018)), therefore we interpret these results to suggest that the 2D fast and 40% CR treatments increased cardiac health relative to 20% CR. These results highlight the complex manner in which the type and magnitude of dietary intervention may improve or degrade cardiac health and may explain the seemingly contradictory results of IF and CR interventions observed in other rodent models (Ahmet *et al*. (2005, 2011)).

Examining the effect of dietary intervention on other aspects of physiological health suggest that the 40% CR treatment was not universally beneficial. The 40% CR group had the lowest hearing ability across the entire auditory range tested, whereas hearing ability was greatest in the AL group (Figure 2). These results contradict previous studies that found caloric restriction prevented age-related hearing loss (Someya *et al*. (2007, 2010)). Similar to hearing ability, we observed bone mineral density was lowest in 40% CR and greatest in AL diet (DX_BOMD; Figure 2). These result were consistent with human clinical trial which showed cardiovascular function was improved and bone mineral density was degraded following a 25% CR intervention (Villareal *et al*. (2006, 2016); Kraus *et al*. (2019)). By measuring multiple aspects of physiology in a large outbred mouse population, we identified contrasting effects of CR and IF on health. With continued observation, we will determine whether the year one effects will have lasting physiological effects on health and explain the physiological mechanisms by which dietary intervention extends lifespan.

### The effect of select genetic variants on physiological health may be as impactful as dietary intervention

The majority of traits (31 of 36) derived from six phenotypic assays exhibited significant genetic heritabilities (Figure 3). Genetic mapping analyses with directly measured and PC derived traits identified both diet-independent and diet-dependent QTLs associated with distinct aspects of physiology (Table 1). We found the effect of founder alleles at some QTLs were as strong or stronger than the effect of dietary intervention. For instance, the difference between positive and negative founder-allele-effects for the lead genotyped variant at the chromosome 5 bone mineral content QTL (0.77, Figure 5F) exceeded the negative effect of 40% CR diet (−0.57). This suggested that the potentially detrimental effect of 40% CR on bone mineral content may be offset by the beneficial effect of the PWK and WSB founder alleles. Similarly, the negative effect of 40% CR on hearing ability (−0.31) could be offset by the significantly positive effect of the WSB, PWK, CAST alleles (1.19) at the chromosome 10 QTL (Table 1). The candidate genes at these loci maybe fruitful targets for genetic manipulation or therapeutic intervention to either mimic beneficial or ameliorate detrimental effects of caloric restriction and intermittent fasting. Finally, the extensive genetic correlations identified between traits, both within and between clusters, suggests that interventions may have pleiotropic effects (perhaps positive or negative) beyond the focal trait.

### Cardiac morphology and function is shaped by diet-dependent genetic associations

Cardiac morphology and function were the only physiological traits for which we identified significant diet-dependent QTLs. Variation in cardiac pumping efficiency, quantified with PC_ECE2, was associated with an AJ specific allele that increased function in 20% CR, 1D, and 2D fast treatments and decreased function in the AL and 40% CR treatments (Figure 6D). Interestingly, the diet-dependent effect of the NZO allele at this locus was nearly opposite that of AJ and the difference between these alleles in the AL (0.500) and 2D fast (0.746) treatments was of similar magnitude of the difference between diets (0.631). We highlight this example to illustrate that the beneficial or detrimental effects of diet maybe ameliorated by genetic variants segregating within the DO mouse population. These results provide additional support for the hypothesis that cardiac efficiency maybe altered to the same degree as CR or IF with genetic manipulation or therapeutic intervention to phenocopy the AJ or NZO allele. Additionally, the large diet-specific effects of the 129 and NZO alleles (Figure 6E,F) suggest that similar approach could be utilized to manipulate LV posterior wall thickness. The decline in cardiac health in response to diet and age is a leading risk factor for reduced lifespan in human populations (Dwyer-Lindgren *et al*. (2016)). These results clearly demonstrate that functional variants are segregating within the DO population to modulate cardiac morphology and function in a diet-specific manner and suggest possible interventions to protect against the diet-induced or age-related decline of cardiac health.

### Future considerations and limitations

In summary, we found that multiple aspects of physiology in early adulthood mice change in response to dietary intervention. Using a diverse set of experimental assays, we identified dietary interventions that may improve or degrade health along multiple axes of physiology. It is unknown how changes observed at one year of age, after six months of treatment, will impact health at later ages. As these mice age, we will continue to monitor them with the ultimate goal of identifying the physiological mechanisms by which dietary interventions improve or deteriorate health at advanced age.

## Supporting information

Standard Operating Procedures

## Acknowledgments

The authors would like to acknowledge Natalie Telis, J. Graham Ruby, Nick van Bruggen and David Botstein for their comments on the manuscript. Funding was provided by Calico Life Sciences LLC.

## Supplemental Material

### Heritability analysis model details

We estimated heritability by fitting the Bayesian model *Y* = *Xβ* + *ϵ* where *ϵ* follows multivariate normal distribution with mean 0 and covariance matrix *σ*^2^(2*h*^2^*K* + (1 − *h*^2^)*I*) where *σ*^2^ is the total phenotypic variance, *h*^2^ is heritability, *K* is the kinship matrix and *I* is identity matrix. The prior information is as follows:

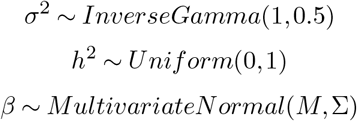

where *M* = [0, 0, 0, 0, 0] and Σ = 2*I*_5*X*5_.

### Genetic correlation analysis model details

Considering two traits *Y*_1_ and *Y*_2_, we estimated genetic correlation by fitting the Bayesian model: 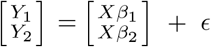 where *ϵ* follows multivariate normal distribution with mean 0 and covariance matrix 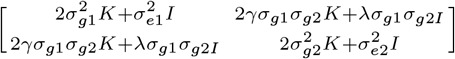 where *K* is the kinship matrix; *I* is the identity matrix; 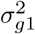 and 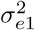 are genetic and environmental variance for trait *Y*_1_ respectively; 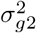 and 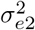 are genetic and environmental variance for trait *Y*_2_ respectively; *γ* is genetic correlation and *λ* represents the correlation due to an individual’s environment. The prior information is as follows:

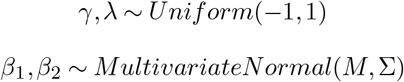

where *M* = [0, 0, 0, 0, 0] and Σ = 2*I*_5*X*5_. 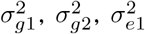 and 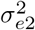 are estimated by fitting each trait individually with diet as fix effect and kinship as random effect using maximum likelihood method.

## Supplemental Tables and Figures

**Table S1.**
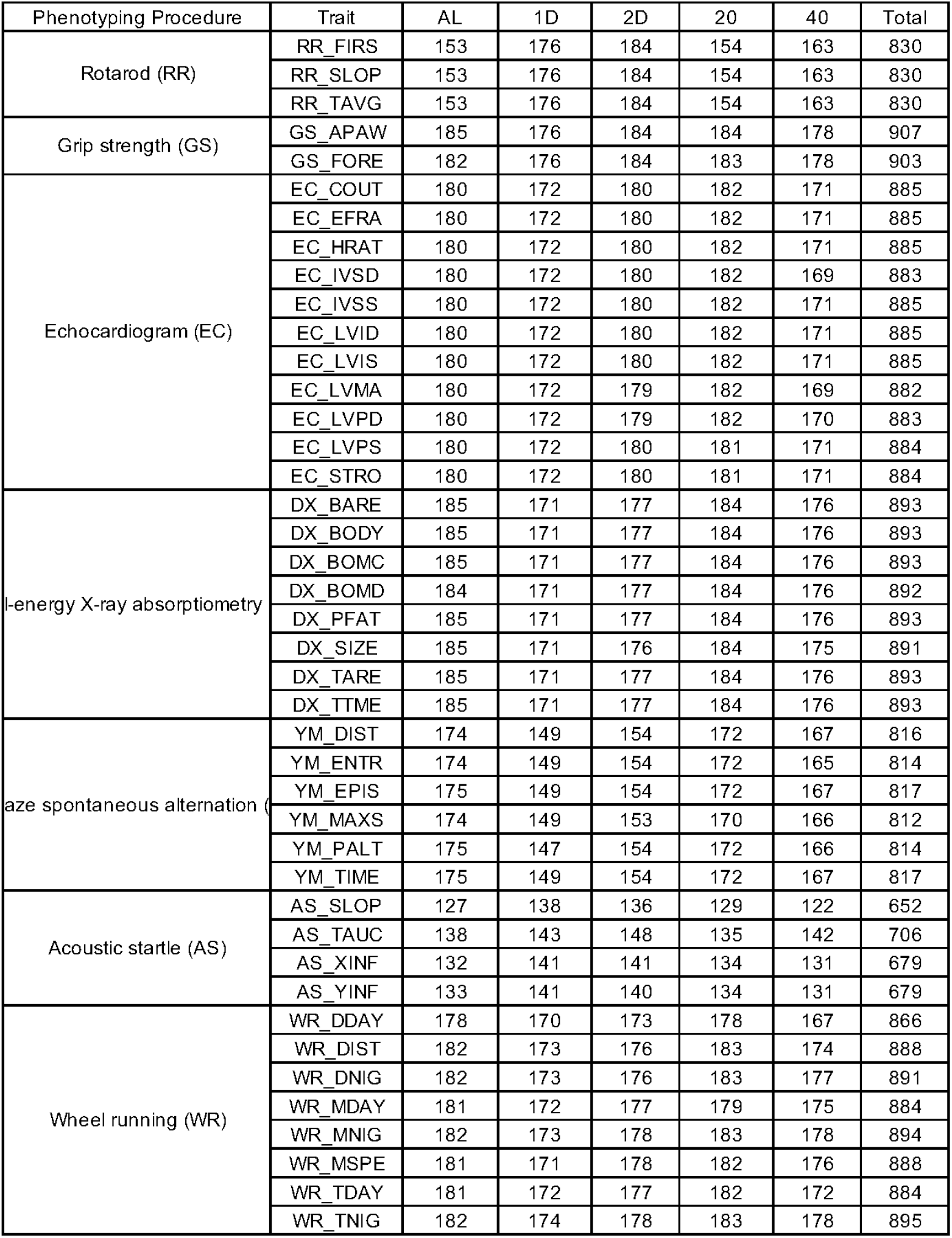
**Total number of samples per trait and per diet after outlier removal**.

**Table S2.** **Significance threshold for unsupervised hierarchical clustering analysis**.

| Cluster size | Threshold |
| --- | --- |
| 2 | 0.55 |
| 3 | 0.59 |
| 4 | 0.64 |
| 5 | 0.66 |
| 6 | 0.68 |
| 7 | 0.71 |
| 8 | 0.71 |
| 9 | 0.75 |
| 10 | 0.73 |
| 11 | 0.75 |
| 12 | 0.75 |
| 13 | 0.76 |
| 14 | 0.77 |
| 15 | 0.79 |

**Table S3.**
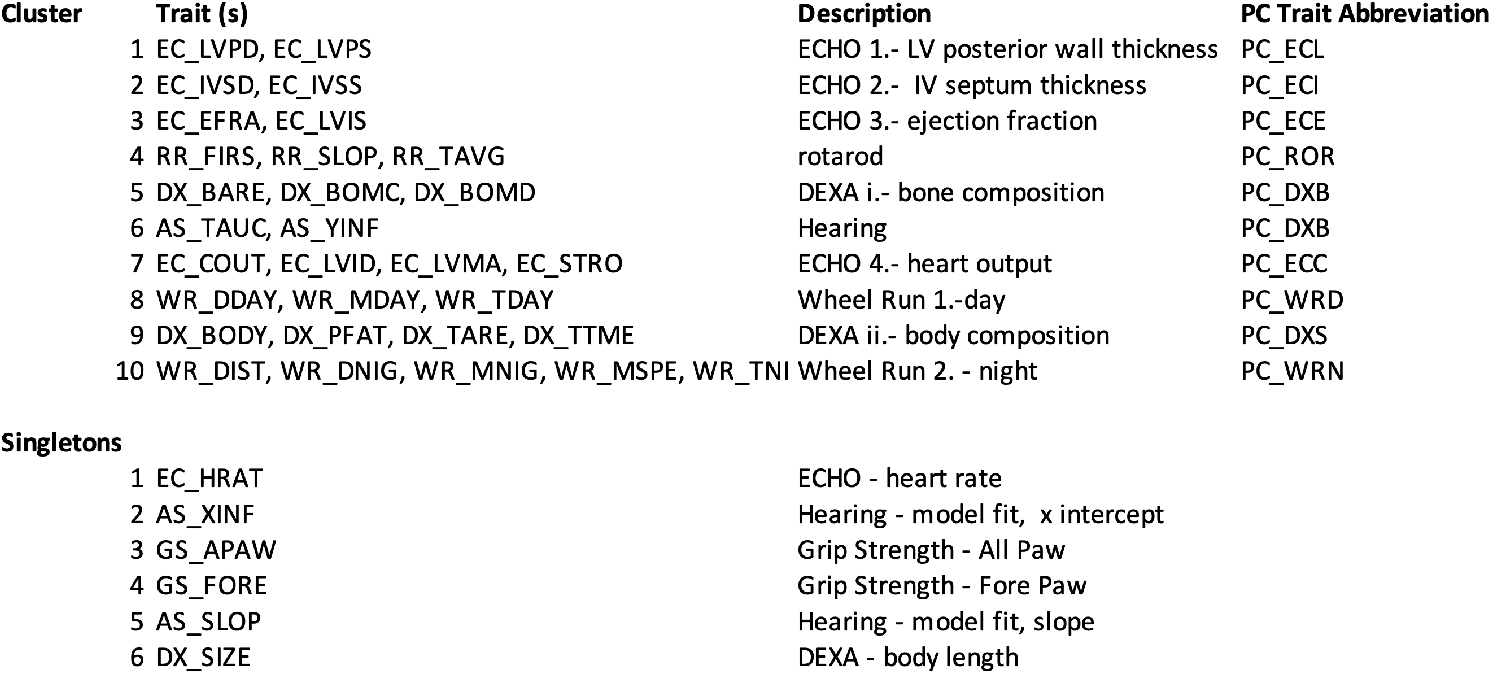
For trait groups identified in hierarchical clustering analysis, we list the directly measured and the principal component derived traits.

**Table S4.** Reduced genotype x diet association model test. For each lead marker at a GxD interaction QTL, we compare the LOD scores of full (Model I) and reduced (Model II) genetic association models. Reduced models test the effect of four, non AL diets in isolation, and for the single diet with the maximum difference between Model I and Model II LOD score, the three possible two diet combinations are also tested.

|  | Diet | Model | LOD_model_I<br>-<br>LOD_model_II |
| --- | --- | --- | --- |
| PC_ECE2<br>Chr16:<br>UNC26651633 | 1D | Model I: $Y = \text{Diet} + G + \text{Diet1D} * G + \text{Diet20} * G + \text{Diet2D} * G + \text{Diet40} * G + \text{Kinship} + E$<br>Model II: $Y = \text{Diet} + G + \text{Diet2D} * G + \text{Diet20} * G + \text{Diet40} * G + \text{Kinship} + E$ | 3.4 |
| | 20 | Model II: $Y = \text{Diet} + G + \text{Diet1D} * G + \text{Diet2D} * G + \text{Diet40} * G + \text{Kinship} + E$ | 7.2 |
| | 2D | Model II: $Y = \text{Diet} + G + \text{Diet1D} * G + \text{Diet20} * G + \text{Diet40} * G + \text{Kinship} + E$ | 7.7 |
| | 40 | Model II: $Y = \text{Diet} + G + \text{Diet1D} * G + \text{Diet20} * G + \text{Diet2D} * G + \text{Kinship} + E$ | 1.8 |
| | 1D/20 | Model II: $Y = \text{Diet} + G + \text{Diet2D} * G + \text{Diet40} * G + \text{Kinship} + E$ | 8.1 |
| | 20/2D | Model II: $Y = \text{Diet} + G + \text{Diet1D} * G + \text{Diet40} * G + \text{Kinship} + E$ | 11.6 |
| | 20/40 | Model II: $Y = \text{Diet} + G + \text{Diet1D} * G + \text{Diet2D} * G + \text{Kinship} + E$ | 8.9 |
| AS_YINF<br>Chr9:<br>UNC16962149 | 1D | Model I: $Y = \text{Diet} + G + \text{Diet1D} * G + \text{Diet20} * G + \text{Diet2D} * G + \text{Diet40} * G + \text{Kinship} + E$<br>Model II: $Y = \text{Diet} + G + \text{Diet2D} * G + \text{Diet20} * G + \text{Diet40} * G + \text{Kinship} + E$ | 3.6 |
| | 20 | Model II: $Y = \text{Diet} + G + \text{Diet1D} * G + \text{Diet2D} * G + \text{Diet40} * G + \text{Kinship} + E$ | 0.7 |
| | 2D | Model II: $Y = \text{Diet} + G + \text{Diet1D} * G + \text{Diet20} * G + \text{Diet40} * G + \text{Kinship} + E$ | 3.9 |
| | 40 | Model II: $Y = \text{Diet} + G + \text{Diet1D} * G + \text{Diet20} * G + \text{Diet2D} * G + \text{Kinship} + E$ | 4.8 |
| | 1D/40 | Model II: $Y = \text{Diet} + G + \text{Diet20} * G + \text{Diet2D} * G + \text{Kinship} + E$ | 8.9 |
| | 20/40 | Model II: $Y = \text{Diet} + G + \text{Diet1D} * G + \text{Diet2D} * G + \text{Kinship} + E$ | 7.3 |
| | 2D/40 | Model II: $Y = \text{Diet} + G + \text{Diet1D} * G + \text{Diet20} * G + \text{Kinship} + E$ | 7.2 |
| PC_ECL1*<br>Chr2:<br>JAX00486864 | 1D | Model I: $Y = \text{Diet} + G + \text{Diet1D} * G + \text{Diet20} * G + \text{Diet2D} * G + \text{Diet40} * G + \text{Kinship} + E$<br>Model II: $Y = \text{Diet} + G + \text{Diet2D} * G + \text{Diet20} * G + \text{Diet40} * G + \text{Kinship} + E$ | 6.2 |
| | 20 | Model II: $Y = \text{Diet} + G + \text{Diet1D} * G + \text{Diet2D} * G + \text{Diet40} * G + \text{Kinship} + E$ | 5.6 |
| | 2D | Model II: $Y = \text{Diet} + G + \text{Diet1D} * G + \text{Diet20} * G + \text{Diet40} * G + \text{Kinship} + E$ | 1.8 |
| | 40 | Model II: $Y = \text{Diet} + G + \text{Diet1D} * G + \text{Diet20} * G + \text{Diet2D} * G + \text{Kinship} + E$ | 1.8 |
| | 1D/20 | Model II: $Y = \text{Diet} + G + \text{Diet2D} * G + \text{Diet40} * G + \text{Kinship} + E$ | 11.9 |
| | 1D/2D | Model II: $Y = \text{Diet} + G + \text{Diet20} * G + \text{Diet40} * G + \text{Kinship} + E$ | 9.4 |
| | 1D/40 | Model II: $Y = \text{Diet} + G + \text{Diet20} * G + \text{Diet2D} * G + \text{Kinship} + E$ | 8.0 |

\*Note: EC\_LVPD (Chr2: UNCHS004526) has the same pattern as PC\_ECL1.

**Fig. S1.**
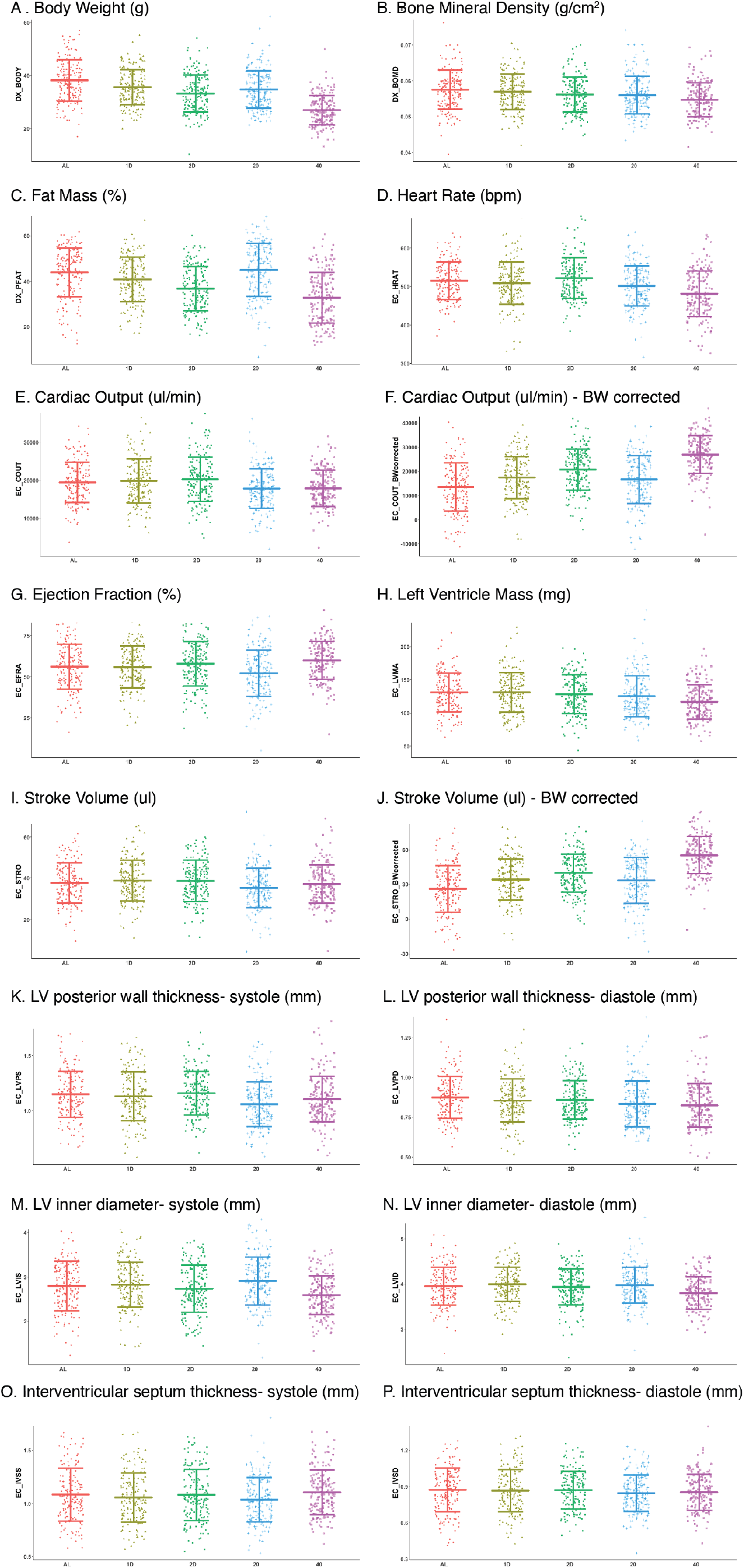
Sixteen DEXA and echocardiogram derived trait values for each diet. Horizontal bars display Mean +/-SD. For cardiac output (EC_COUT) and stroke volume (EC_STRO) we present the raw values and body weight corrected values (calculated following the same procedure as applied to grip strength and rotarod).

**Fig. S2.**
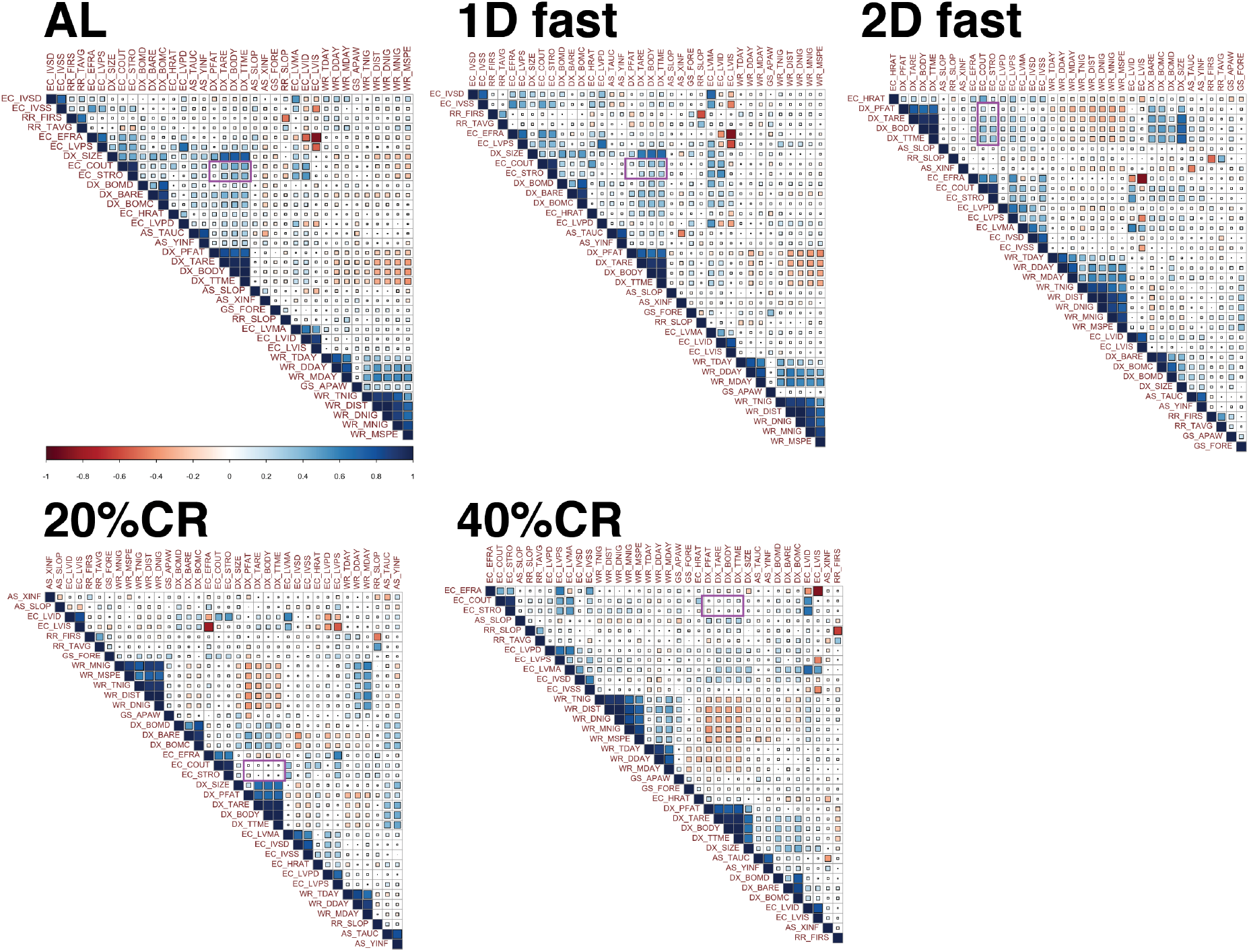
Diet-specific pairwise phenotypic correlation values. Size and color of squares represent the positive (blue) or negative (red) correlation values. Purple box highlights pairwise correlations between cardiac output and stroke volume (EC_COUT, EC_STRO) and multiple body composition traits (DX_PFAT, DX_TARE, DX_BODY, and DX_TTME).

**Fig. S3.**
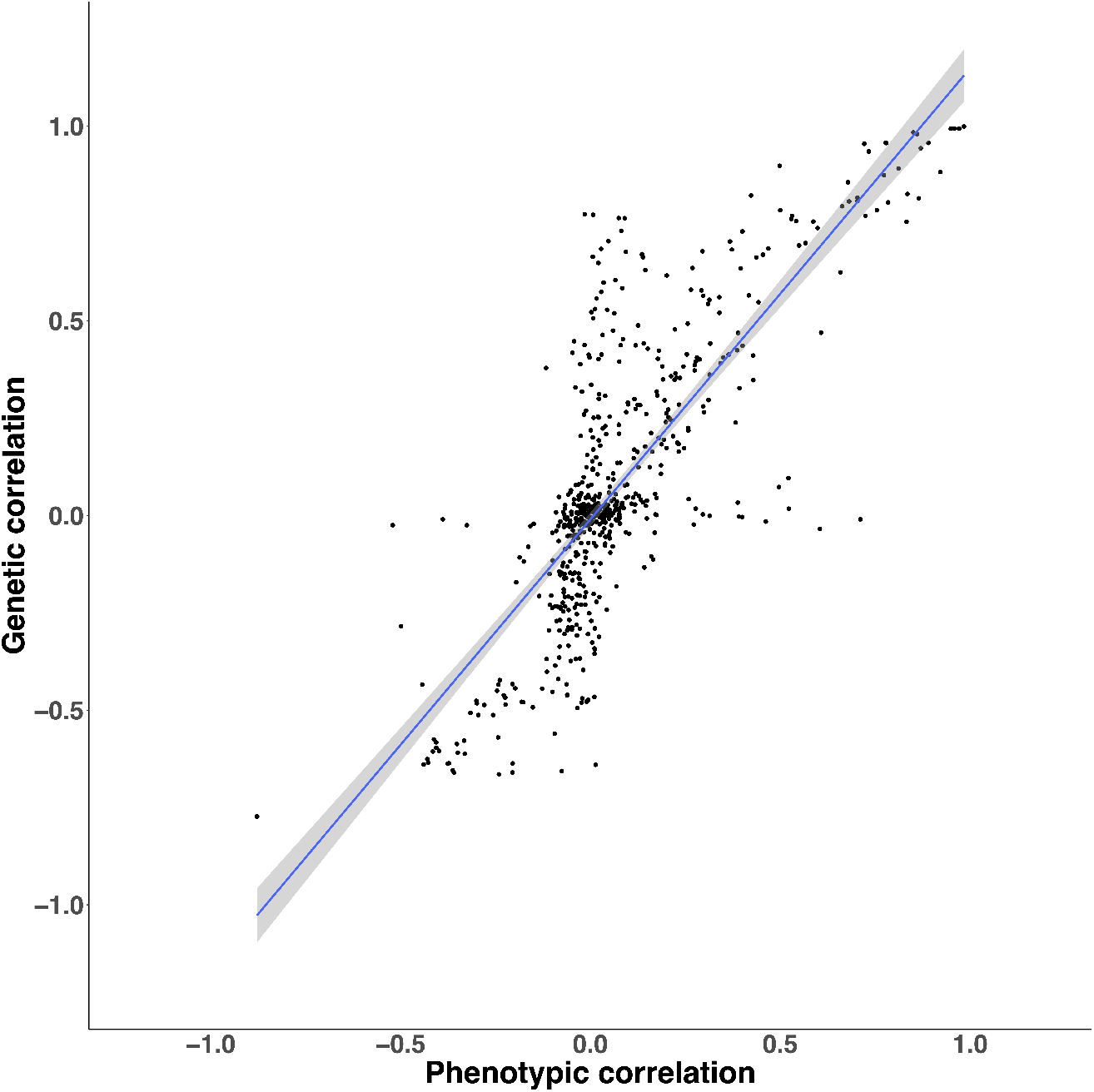
Scatterplot of phenotypic versus genetic correlations. Grey line depicts linear correlation with 95% CI in shaded area.

**Fig. S4.**
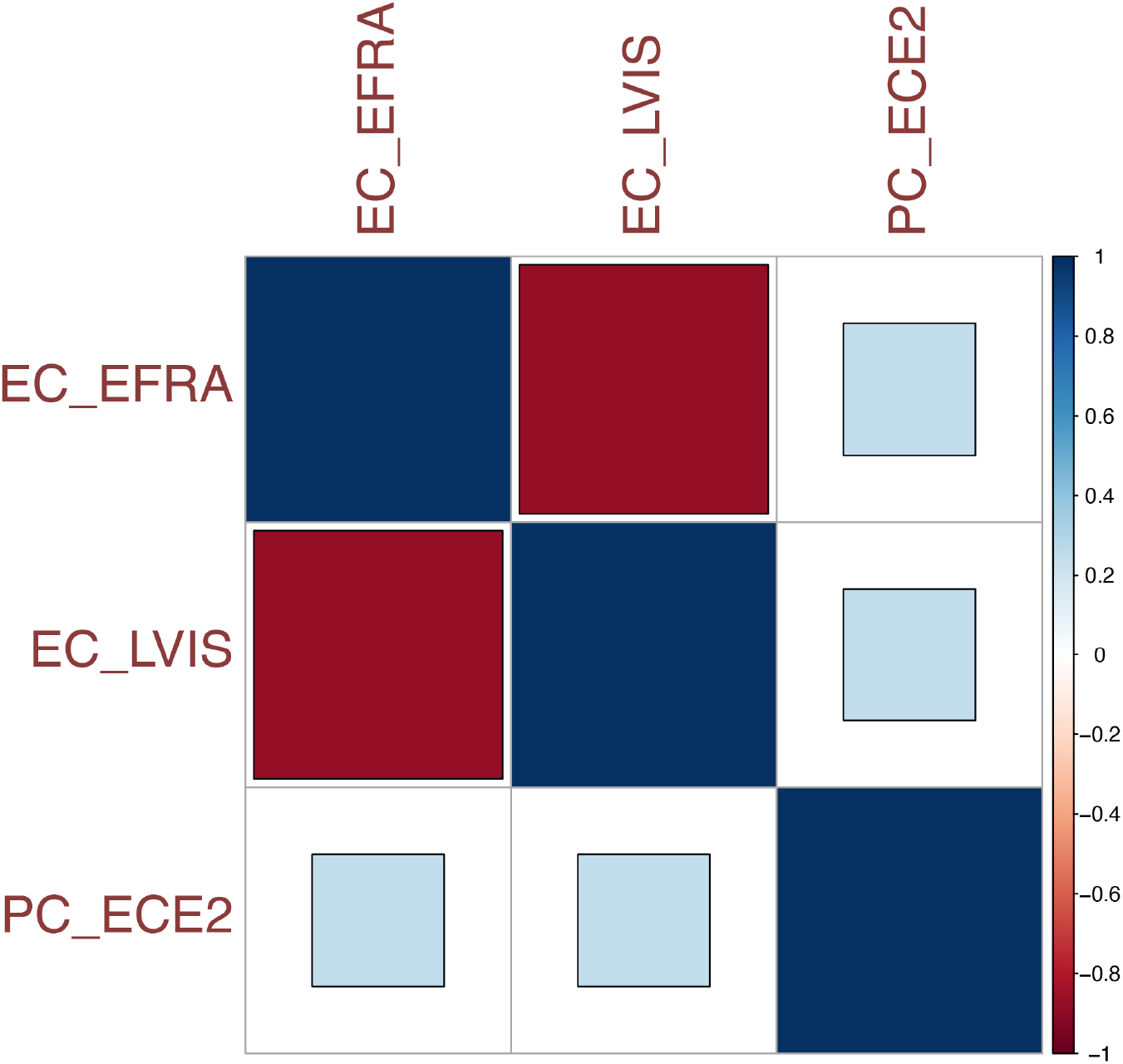
Pairwise Pearson correlation values between PC_ECE2 and the two directly measured traits used to calculate this principle component analysis trait: EC_EFRA and EC_LVIS.

**Fig. S5.**
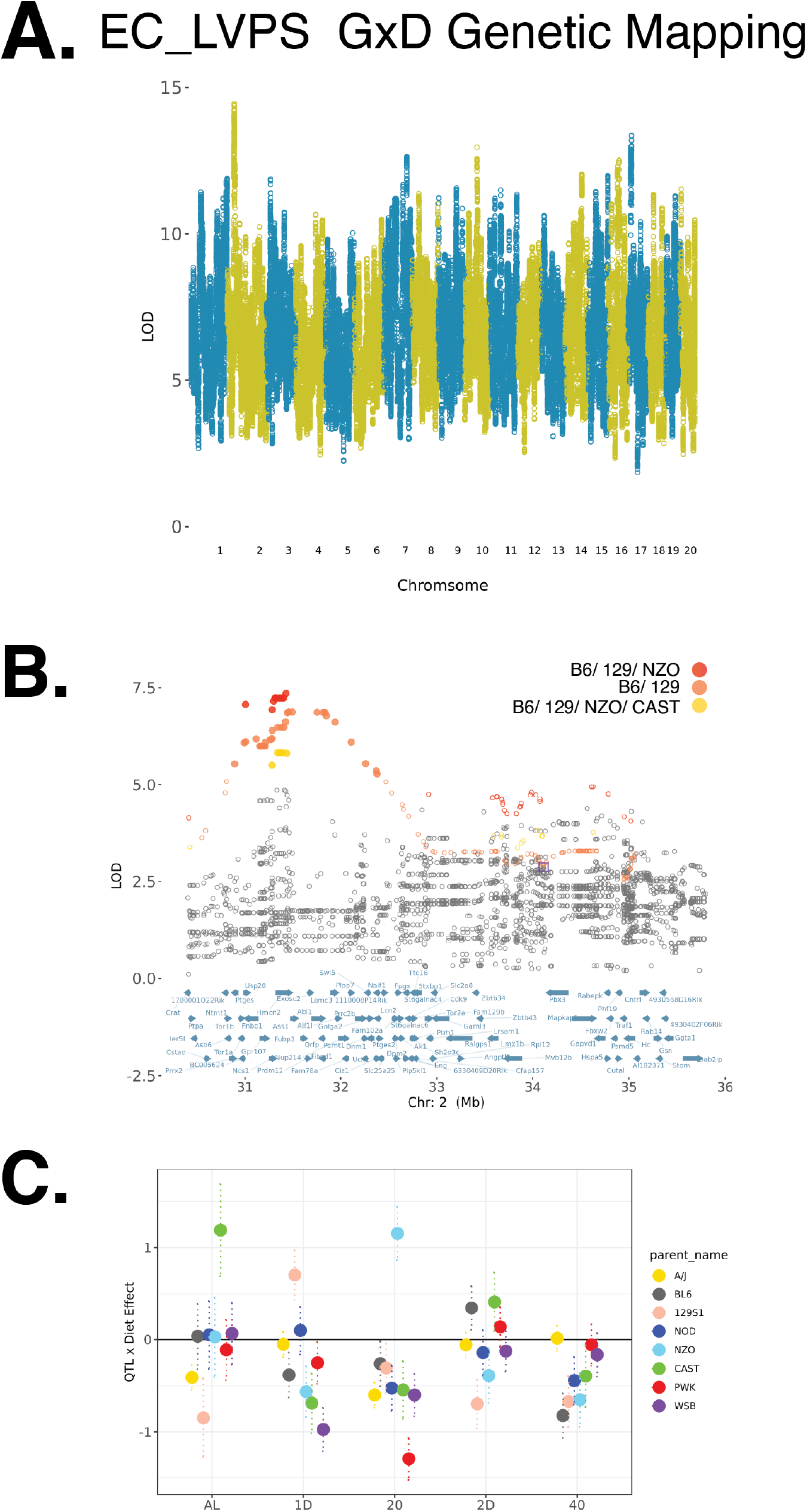
A. Manhattan plot of diet-dependent genome-wide linkage mapping results for EC_LVIS. B. Fine-mapping of chromosome 2 locus. Rank 1, 2, and 3 FAP variants shown in red, orange, and yellow circles. C. Diet-specific effect of lead genotyped variant for each of the eight founder variants.

## Notes

### Competing Interest Statement

The authors have declared no competing interest.

