## Supplementary material for "Intermittent fasting and caloric restriction interact with genetics to shape physiological health in mice": Standard Operating Procedures

**Wheel Running**

#### Laboratory Information

| | |
|---|---|
| Department | The Jackson Aging Center |
| Principle Investigator(s) | Gary Churchill |
| Creation date | January 1, 2015 |
| SOP Author | Laura Robinson |
| Revision date | March 11, 2019 |

#### Brief Description of Procedure of Experiment

Wheel running behavior in mice is a simple and easily quantifiable measure of behavior that can be assessed in the home cage and with little interruption (Sherwin, 1998). In addition to providing enrichment in the home café environment, it is well reported that wheel running behavior is sensitive to genetic alterations (Lightfoot et al 2008) and can also provide phenotypic assessment for circadian activity (reviewed in Pendergrast et al 2014) It has also been vital in demonstrating subtle phenotypes in neuromuscular and neurodegenerative models that may not be readily detectable in motor based behavioral tasks (ie; treadmill, open field) (Mandillo et al 2014) and may even implicate motivational behavior phenotypes (Rhodes et al 2005; Coyle et al 2008).

#### Hazard Assessment-Equipment/Mechanical/Electrical Hazard

NA

#### Hazard Mitigation

NA

#### Procedure

The apparatus used is the *Med Associates* low profile running wheels with a wireless transmitter will be used for these studies (Figure1). The wheels are similar to the types of running wheels currently used within the Research Animal Facility (RAF) for enrichment with the exception that they contain the wireless transmitter in the base. This is a high throughput system that can accommodate up to 120 running wheels at one time (per computer).

Figure 1.

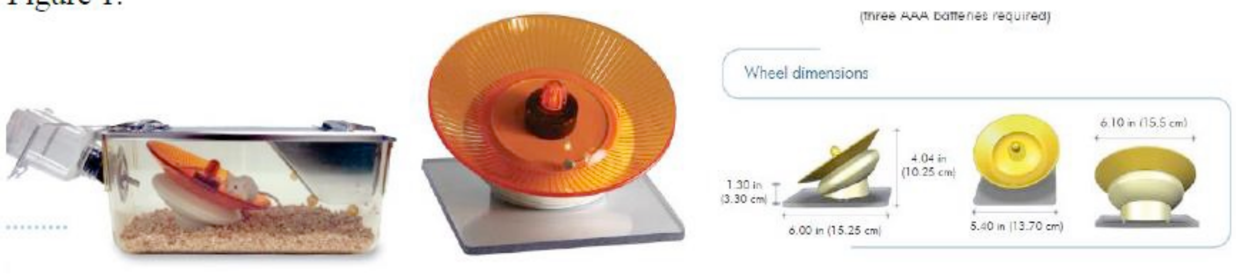

Given the dimensions of wheels, they will not fit in to the standard duplex cage (which is too narrow) or the standard wean cages (which has a limited ceiling height with the wire bar cage top). Therefore, the best suited cage to accommodate the wheels is the Innovive disposable caging or a shoebox cage with a modified wire bar lid that eliminates a food hopper. It is important to note that these cages have a food hopper which sits in the center of the cage which presents an obstruction; therefore the hopper will be removed. Food will be provided on the cage floor. Cages with food on the floor will be changed to a new cage on a weekly basis.

Mice will be individually housed during these studies in order to get accurate accounts of individual mice. A typical study lasts from 5-10 days. However, depending on the objectives of the experiment, mice may be housed with the wheel for up to 30-90 days. These longer term studies are typically needed to assess circadian pattern phenotypes and may include phase shifting studies. In the case of phase shifting studies, mice will first be acclimated to the running wheels on the standard 12:12 L:D period (entrainment). Following a 5-10 day period at standard L:D, mice may be shifted to lights going off earlier (by 1-2 hours) or later (by 1-2 hours) or moved to a D:D schedule (24 hours dark) at which time phase shifting and phase delays can be evaluated.

**2. Anesthetic/Analgesic Regimen**

**NA**

**3. Post Procedure Care**

No adverse outcomes are anticipated. Animals will be monitored for normal activity and appearance as part of daily cage checks. Normal daily husbandry activities can occur including daily cage checks, replenishing food hoppers and water, bedding changes, and health monitoring. Investigators will be responsible for regular weekly cage changes. It will be critical to identify consistent periods within the day that staff can enter the rooms. During DD periods, staff will be trained to use red lights and/ or headlamps with red light.

At the conclusion of the testing periods, mice will return to normal housing in the housing room and the equipment can be sanitized thoroughly by CMQ approved sanitization agents (e.g., Virkon).

CLAM will be contacted if any abnormal or unexpected behaviors are observed.

JACKSON AGING CENTER

### Rotarod

#### Laboratory Information

| | |
|---|---|
| Department | The Jackson Aging Center |
| Principle Investigator(s) | Gary Churchill |
| Creation date | January 1, 2015 |
| SOP Author | Laura Robinson |

#### 1. Procedure Description

The Rotarod task is a classic test of motor coordination which is used to assess either motor/coordination phenotypes in mice or the effects of test compounds.
`

The apparatus used is the *A Ugo-Basile rotarod* (Figure1) consisting of five drums (3 cm diameter), suitably machined to provide grip for the mice will be used for these studies. Six flanges divide five 5.7cm lanes, enabling five mice to be evaluated simultaneously. When a mouse falls off its cylinder section on to the plate below, the plate trips thereby recording the animal's endurance time in seconds. The height to fall is recorded at 16cm

Figure 1. (From www.ugo-basile.com)

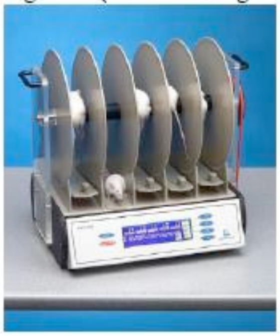

For all tests, mice are acclimated to the testing room at least 60 minutes prior to testing.

There are two separate protocols depending on phenotyping or drug compound testing:

1. Phenotyping:

   For a typical single test, the protocol is as follows: At the beginning of the test session, the mouse is placed on a fixed speed rod (rotating at 4 rpm) that increases linearly to a maximum rpm of 40, over 300 seconds. Mice are given 3 successive trials (approximately 1 minute inter-trial interval to clean the rod between trials).

   For Longitudinal testing, as often in the case of progressive neurodevelopmental models (e.g, Huntington's R6/2mice) a longer protocol may be necessary since normal healthy mice (WT) quickly learn this test and retesting procedures a ceiling effect. In these cases, a 10 minute(min.) protocol will be used such that the ramp will increase from 4-40 RPM over 600 seconds with 3 consecutive trials as above.

2. Effects of Test Compounds:

This experimental design is used to determine if a therapeutic agent produces motor impairing effects relative to vehicle treated controls. For this test, mice will initially be trained in the fixed speed paradigm (4rpm for 3 consecutive 1 min. trials). This is to ensure that all subjects demonstrate identical baselines (for inclusion in this test) prior to being administered test compound. At the conclusion of the training session, test subjects will be administered test compound or vehicle control by various routes of administration including: CMQ93-25 subcutaneous (s.c.), LAH07-01 oral (p.o.), in accordance with the following IACUC approved routine procedures (CMQ93-25 and LAH07-01, respectively); with dose volumes of 10-20 ml/kg. Following the pre-determined pretreatment time (as determined by drug pharmacokinetics, typically 0-60 min.), subjects will be placed onto the rod at a starting speed of 4rpm which will increase linearly to 40RPM over 600 seconds. For these tests, 3 trials will be used and separated by 30 or 60 min. rest periods (i.e, 30 min. post, 60 min. post, 90 min. post). The purpose of separating the trials by 30 or 60 min. is to assess the half-life of drug on the behavioral effects (when the drug starts to lose efficacy).

At the conclusion of testing, mice are returned to their home cages. Between subjects, the rods, lanes, and floor of the rod are cleaned with 70% ETOH and additional sanitization with CMQ approved agents (e.g., Virkon) will be completed at the end of the testing session.

#### 2. Anesthetic/Analgesic Regimen

## NA

#### 3. Post Procedure Care

No adverse outcomes are anticipated. Drug effects are transient with behavioral effects typically attenuating by the end of the test session. Animals will be monitored for normal activity and appearance after being returned to the home cage and daily thereafter for food and water intake and grooming as part of daily cage checks. CLAM will be contacted if any abnormal or unexpected behaviors are observed.

Research Laboratory Standard Operating Procedure

### Grip

**Laboratory Information**

| | |
|---|---|
| Department | The Jackson Aging Center |
| Principle Investigator(s) | Gary Churchill |
| Creation date | January 1, 2015 |
| SOP Author | Laura Robinson |
| Revision date | March 26, 2019 |

#### Brief Description of Procedure or Experiment

A commercially available Grip-strength meter (Bioseb) is used to measure forelimb grip strength and/or combined forelimb/hindlimb grasp strength as an indicator of neuromuscular function, which takes advantage of the mouse's natural behavior of grasping. For all measurements, a wire grid coupled to a strain gauge which measures peak force (g) is used. The mouse is lowered gently towards the wire grid and instinctively grasps the bar with its paws. Care is taken to ensure the mouse is gripping the grid properly, with both front paws only (for forepaw measurements) or with all four paws (for the combined measurements). Once an appropriate grip is assumed, the animal is gently and firmly pulled from the grid until it releases its grasp. Peak force is measured in g for a total of 6 consecutive trials (typically 3 consecutive forepaw followed immediately by 3 consecutive all four paws) and an average is calculated for each animal**.**

#### Hazard Assessment - Equipment/ Mechanical / Electrical Hazard

NA

#### Hazard Mitigation

NA

#### Procedure

Before Test:

- Weigh and label mice – record weights in new  grip template file
- Arrange boxes of mice to be tested on rolling cart and move cart adjacent to grip testing area to habituate mice for 1 hour.

Materials:

- Setup weaning cage with no shavings and wire cover to work as a transition cage.
- Attach grid to grip meter using thumb screw
- Set BIOSEB grip meter on a solid table with meter base against a wall for additional stability
- Set laptop beside meter
- Plug USB software key into USB port on the laptop
- Attach cable from Grip Meter to another USB port on laptop computer

BIOSEB Grip Meter	Grip Meter to Laptop	Transition Cage

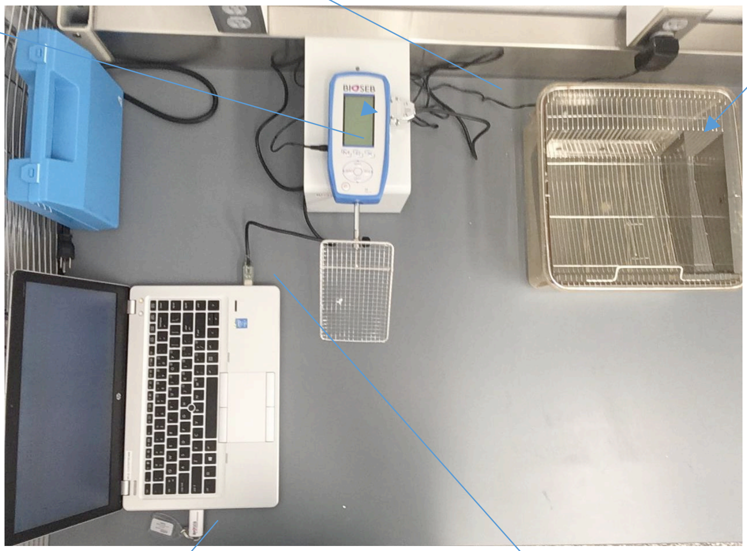

USB Software Key	Grip Meter to Laptop Cable

- Turn on grip meter by pressing power button

- Select the MAX button to proceed
- Set up internal memory to store a series of measures by pressing the M button
- Press TDX button to select "STATS"
- Check that the measures/samples value is set to desired samples per mouse ('6' in the 3 forepaw and 3 all paw), the operator is 1, and the unit is in grams (g). To adjust any of these values, press the TDX button and increase or decrease with Up and Down arrows on panel. Press TDX again to store change and X button to save all changes
- At the bottom of the main screen you will see STAT 001/1. The first number (001) represents the trial; the second (1) represents the number of samples within that trial. 001/3 would be the first trial and third sample in a series, or first test mouse on its third test/pull
- Open the BIO-CIS software from the laptop's desktop and select Connect Device (upper left)
- Click OK to accept settings
- Select Start Protocol. The Device drop down should read 'Grip Test.' In the Protocol drop down, select the '3 Forepaw 3 All Paw' protocol
- Click OK and an excel window will open with accompanying Data Acquisition pop-up window

Note: The internal memory will save up to 100 measures, at that point a FULL MEMORY message will appear on the screen. Since there are 6 samples per mouse as in this protocol, then 16 mice can be tested before memory must be dumped and cleared (16 mice * 6 measures = 96 measures)

Test:

- Wipe work surface with 70% ethanol so first animal will experience same environment as subsequent animal
- Retrieve first mouse to be tested by gently grabbing mid-tail
- Press ZERO button to tare meter, anything within the +0.0-1.0 range is acceptable since the meter only records the Max Force value
- Gripping mouse gently mid-tail, suspend just above grip grid/bar
- Wait for a slight reaching or grabbing response toward the bar then let the mouse grab grid, gently lower hand holding tail a little bit, and pull back horizontally to the table with a constant force

Note: When using the grid make sure the mouse has grasped somewhere between the middle and T-bar portion of the grid.

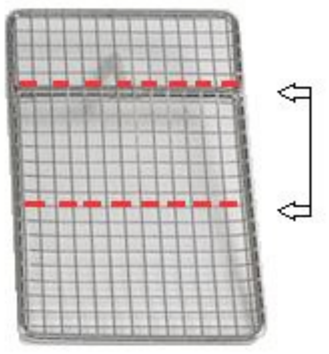

- After mouse has released bar/grid, the meter will display the max grip force in grams. Press TDX button to accept and record this value in the internal memory or ZERO again if it was a faulty test (i.e. mouse didn't grip properly)

Let mouse rest briefly on table after pull then repeat until all 6 samples for that mouse has been reached.

- Following each accepted test, the STAT x/x values will increase incrementally to help you keep track of trials and samples (For a 3 sample setup with 2 mice: 001/1, 001/2, 001/3, 001/4, 001/5, 001/6, 002/1, 002/2, 002/3 etc.)
- After mouse has been tested, deposit in Transition Cage
- Wipe down table with 70% ethanol to maintain a clean working surface.
    - Do not press on, or clean grid while it is attached to the meter**.**
- Retrieve next mouse and repeat until 16 mice have been tested.
- Since BIOSEB Grip Meter memory is now nearly full it is necessary to transfer that data into the excel file on the laptop. In the Data Acquisition pop-up window on the laptop, click Read Value. All data from the grip meter's memory will auto populate in the excel file. Save data in prepared grip template.
- Once data values have been transferred and saved, reset the meter's memory

  a. Press M button.

  b. Press ZERO button.

  c. Press M button to erase all data.

  d. Once reset the STAT value should read 001/1 once more

- Continue testing the next set of 16 mice until all mice are tested.
- When testing is complete, reset memory and power off meter
- Clean Transition Cage with 70% ethanol between boxes of mice to maintain a clean area
- Remove grid from meter, clean with 70% ethanol, and store beside meter (to prevent any possible damage if meter gets knocked)

#### Anesthetic/Analgesic Regimen

NA

#### Post Procedure Care

No adverse outcomes are anticipated. Animals will be monitored for normal activity and appearance after being returned to the home cage and daily thereafter for food and water intake and grooming as part of daily cage checks.

#### References (if applicable)

Meyer, O.A., Tilson, H.A., Byrd, W.C., Riley, M.T. 1979. A method for the routine assessment of fore and hindlimb grip strength of rats and mice. Neurobehav. Toxicol.. 1: 233-236

Research Laboratory Standard Operating Procedure

**Ultrasonography, including Echocardiography**

**Laboratory Information**

| | |
|---|---|
| Department | The Jackson Aging Center |
| Principle Investigator(s) | Gary Churchill |
| Creation date | January 1, 2015 |
| SOP Author | Laura Robinson |
| Revision date | March 26, 2019 |

**Brief Description of Procedure or Experiment**

During Echocardiography, the mouse is anesthetized using Isoflurane. Once anesthetized, the fur in the area necessary to achieve an echocardiograph is removed using either clippers or a depilatory agent, such as “Nair”. Heated ultrasound transmission gel is applied to the animal’s skin, the ultrasound probe is brought to contact with the animal’s skin to measure blood flow rates and volumes. The mouse is then wiped clean of ultrasound gel and allowed to recover from anesthesia in a heated environment.

**Hazard Assessment - Equipment/ Mechanical / Electrical Hazard**

**N/A**

**Hazard Mitigation**

**N/A**

**Procedure**

Ultrasonography will be performed using a VisualSonics Inc. (VSI) Vevo 770/2100 high-frequency ultrasound with 30 and 40 MHz probes. The mice will be anesthetized with Isoflurane at concentrations up to 5% and flow rate of 0.8 - 2.0 L/min in oxygen. Ophthalmic ointment is placed on the eyes to prevent drying of the cornea while the mouse is anesthetized and tested. After anesthetic induction, the animals will be placed on a thermostatically-controlled heated platform where isoflurane anesthesia will be maintained by delivery through a close fitting face-mask.  The delivered isoflurane concentration will be the minimal necessary to keep the animals immobile for the duration of the examination; generally this will be between 0.5 and 2%.  During the examination, the animal's heart rate will be monitored through use of an electrocardiograph.  Once in position, the appropriate area of the animal's skin overlying the area of interest will be cleared of hair either by the use of clippers or a depilatory agent, such as "Nair".  If a depilatory is used, it will be allowed to remain in contact with the animal's skin for as brief a time as possible to achieve the desired effect.  After this time, remaining agent will be wiped off and the skin rinsed with water to remove any trace amounts.

Once prepared in this way, heated ultrasound transmission gel will be applied to the animal's skin to act as a coupling-medium.  The ultrasound probe is then brought into contact with the animal's skin through the gel and appropriate images acquired according to the experimental design.  Echocardiography uses pulsed Doppler sonography, applied through the ultrasound probe, to measure blood flow rates and volumes.  At completion of the examination, the transmission gel will be removed by gently wiping the animal down, after which the mice are allowed to recover from anesthesia in a heated environment (either with a heating pad and/or a heat lamp) until ambulatory. Once capable of movement, they will be returned to their normal housing.

When contrast agents are required, PolySon microbubbles (Miltenyii Biotech, Inc.) will be introduced into the vascular stream via tail vein injection (LAH93-25) or through a jugular catheter (LAH02-01). Commercially available microbubbles are composed of a perfluorocarbon gas core encapsulated by a lipid shell, stabilized and shielded by a layer of polyethylene glycol, suspended in normal saline or PBS. The microbubbles are generally 0.5-5um in diameter and therefore do not leave the vascular space. Of microbubbles into the lateral tail vein using a 35-30 gauge(G) needle.  If necessary, contrast agent will be delivered through the previously placed jugular

Animals are anesthetized and prepared as described above except that they are anesthetized with isoflurane diluted in 30% oxygen rather than 100% oxygen.  High concentrations of oxygen will reduce the half-life of circulating microbubbles.  Once prepared in this way, the animal will be injected with a slow bolus of 50uL of microbubbles into the lateral tail vein using a 35-30G needle.  If necessary, contrast agent will be delivered through the previously placed jugular catheter.  Bubbles are cleared between 10-20 minutes and if there is successful delivery then no more than 2 injections per animal will be necessary.  A maximum of 4 injections may occur per one mouse, with no more than 200uL of microbubbles injected.  Bubbles will be re-injected once they have cleared from the system and the image has returned to baseline.

For imaging the colon, mice will receive a corn oil enema (~ 1.0-1.5 ml) as a negative contrast agent, as in Pickhardt et al, 2005, after anesthesia.

Imaging and micro-injecting neonates is also possible using isoflurane or deep hypothermia, also referred to as cryoanesthesia. By using the ultrasound and rail system to image, the user can more accurately inject into the desired sites (heart, brain, etc).

**Anesthetic/Analgesic Regimen**

a. Please list all anesthetics/analgesics used in this procedure in the following table.
   If not applicable, please check here ☐ NA

| Anesthetic Agent | Diluents Used | Dose & Route of Administration (e.g. 1mg/kg I.V.) | Volume |
|---|---|---|---|
| Isoflurane | Oxygen | By inhalation to effect: 1-5% | 1-5% @ 1.5L/min |

**Post Procedure Care**

After imaging, mice are placed in a recovery box. Half of the box is warmed to 80-86°F and animal has the opportunity to move away from the heat source. To avoid overheating, the temperature inside the cage is monitored with a thermometer at rodent level prior to animal placement, and frequently thereafter. When the animals have fully recovered, they are returned to cages and checked once again before the end of the day.

**References (if applicable)**

Pickhardt PJ, Halberg RB, Taylor AJ, Durkee BY, Fine J, Lee FT Jr and Weichert JP. Microcomputed tomography for polyp detection in an in vivo mouse tumor model. PNAS (2005), 102 (9), 3419-22.

Research Laboratory Standard Operating Procedure

### Acoustic Startle Response

**Laboratory Information**

| | |
|---|---|
| Department | The Jackson Aging Center |
| Principle Investigator(s) | Gary Churchill |
| Creation date | January 1, 2015 |
| SOP Author | Laura Robinson |
| Revision date | March 11, 2019 |

#### Brief Description of Procedure or Experiment

When a loud, abrupt and unexpected sound is heard, it is reflex in all mammals to startle. Startle response consists of an abrupt and involuntary extension and flexion of a series of muscles. Although the startle response is a reflex, it can be altered by a variety of stimuli. Startle response is measured in rodents using automated startle chambers, in which a mouse is placed in a clear, acrylic tube attached to a highly sensitive platform that is calibrated to track their startle reflex while being exposed to a series of stimuli at varying decibels and times. Mice are initially exposed to white noise from an overhead speaker, which transitions to a series of randomized, computer generated stimuli ranging in decibel from 70-120db at 40 msec in duration and an interval of 9-22 seconds. The test runs for approximately 30 minutes. Once the test has completed, the mice are removed and the chambers are cleaned for the next cohort of mice.

#### Hazard Assessment - Equipment/ Mechanical / Electrical Hazard

Acoustic Startle testing room emits sounds at varying decibels that could be damaging to the ears.

#### Hazard Mitigation

In order to avoid hearing damage, a person performing this assay should remain in the anteroom during the testing period. They should not enter the testing room until they are certain that testing has completed.

#### Procedure

**Habituation:**

Before transporting animals to the testing room, number the tails and weigh each animal. Load pens onto a rack and transport to the testing room. Let mice habituate in the testing room for one hour. During this time, complete an empty run to ensure all test subjects are evenly exposed to stimuli in adjacent holding area.

**Calibration**

1. Attach and plug in standardization unit into chamber 1. Let the unit warm up for 15 minutes before moving onto the next step.
2. Open the SR-LAB program from the desktop. Under *Run*, select *Diagnostics*.
3. Select *File* in the upper left hand corner of the screen and open the diagnostic database. Open the *Calibrate MDB* file.
4. On the table on the right hand of the screen, select *Remove all* if old calibration data is present.
5. Select the chamber to be calibrated and hit start.
6. Observe the average millivolt value in the bottom right hand corner of the screen and wait until it levels out to a constant value. You will want a reading of 700-710 mV for>10 trials. To adjust, turn the response adjust dial on the right side of chamber.
7. Press stop button to end calibration and remove all to clear data from table.
8. Repeat steps 8-10 for all chambers then close out of the Diagnostics window.

**Setting up Experiment**

9. Mice are tested in acoustic startle chambers with chamber light off and fan on. Make sure to check that chamber switches are set accordingly.
10. In the SR-Lab program, under *File*, select *Study Database*. Always use the most recent database.
11. There are 8 chambers so create as many sessions as needed to test your entire cohort. To create test sessions
    a. Select *File* and *Create Session*
    b. Select *Add Session ID* and name session.
    c. Place a checkmark next to the session definition
        a. “Startle response curve_07sept2016”
    d. Click *Save Session* and either add another session ID or close window.
12. Next, assign subjects to a session:

a) Select *File* and *Create Subjects*.
b) Select session to assign subjects to.
c) Enter chamber number and subject ID.
d) Click *Save* and *Close*.
e) Create an empty session (with no animals) and run this during the habituation period.

**Running Tests**

13. Move the 8 animals that are to be tested from the habituation rack onto a rolling metro cart and bring into testing room. All animals housed in the same box must be tested at the same time. Do not separate mice from a box to be tested during different runs.
14. Place each mouse into pre-assigned chamber inside of the animal enclosure tube and secure with snap-in doors
15. Select *Run* and *Session*. Under *Session ID*, scroll and select your pre-named session. Check all chambers that will be in use.
16. Click *Star*t to begin.
17. A popup message will notify you when the session had ended (this takes about thirty minutes). The session data is automatically saved.
18. Return mice to their home cage and clean the load cell platform with 70% ethanol. Soak paper towels in 70% ethanol and push through the enclosure to thoroughly clean.
19. Repeat steps 13-18 until all mice have been tested.
20. After all mice have been tested and chambers cleaned, wipe enclosures down with Virkon cleaner.

#### Anesthetic/Analgesic Regimen

For phenotyping, no anesthetic or analgesic is administered.

#### Post Procedure Care

There is no required post procedure care for Acoustic Startle Response testing.

#### References (if applicable)

The Jackson Laboratory, Routine Procedure Title:MNB14-02 PRE-PULSE INHIBITION OF THE ACOUSTIC STARTLE REFLEX (PPI). Rizzo, S. 2/20/2017.

Research Laboratory Standard Operating Procedure

### PIXI (In Vivo Bone Densitometry and Body Composition – DEXA)

**Laboratory Information**

| | |
|---|---|
| Department | The Jackson Aging Center |
| Principle Investigator(s) | Gary Churchill |
| Creation date | January 1, 2015 |
| SOP Author | Laura Robinson |
| Revision date | March 19, 2019 |

#### Brief Description of Procedure or Experiment

This test measures skeletal and soft tissue mass, enabling assessment of skeletal and body composition in living or dead mice. It utilizes 2 X-ray sources of separate energy levels, which are absorbed differentially by tissue. The ratio of absorption differences is proportional to bone density and body composition (including fat and muscle tissue). The LUNAR PIXImus II densitometer is used to collect PIXI data. Users must be trained in X-ray safety before use, and are required to wear a radiation safety monitoring badge during use. Because of the special training needed, technicians from the Center for Biometric Analysis perform the testing.

#### Hazard Assessment - Equipment/ Mechanical / Electrical Hazard

Due to the use of X-Ray equipment, there is a risk of exposure to radiation.

#### Hazard Mitigation

In order to avoid radiation exposure, all users must be trained in X-Ray safety before using the PIXImus, and wear a radiation safety monitoring badge.

#### Procedure

Mice to be scanned dead are euthanized with $CO_2$ inhalation. Those scanned alive are anesthetized with Tribromoethanol (0.2ml/10g IP) or with Isoflurane (induced with concentrations up to 5% and flow rate of 0.8 – 2.0 L/min in oxygen and maintained via nose cone between 0.5 and 2%). Ophthalmic ointment is placed on the eyes to prevent drying of the cornea while the mouse is anesthetized and tested. Mice are individually placed on a disposable plastic tray that is then placed onto the exposure platform of the PIXImus. The process to acquire a single scan lasts approximately 4 minutes; data can be manipulated subsequently to obtain specific regions of interest. At the end of the scanning time, mice are removed, recovered from anesthesia, and then returned to their home cage.

In the event that longitudinal studies of body composition are required with individual mice, it is possible that a given mouse will be subjected to the above procedure as many as 6 times at an interval as short as every 5-7 days in some cases. Long-term studies could be up to the entire life of the mouse, with scans conducted at intervals of one month or longer.

#### Anesthetic/Analgesic Regimen

| Anesthetic Agent | Diluents Used | Dose & Route of Administration (e.g. 1mg/kg I.V.) | Volume |
|---|---|---|---|
| Isoflurane | Oxygen | Inhalation to effect ~2% | |
| Tribromoethanol | Sterile PBS | 400 mg/kg IP | 0.2 ml/10 g body weight |

| Anesthetic Agent | Diluents Used | Dose & Route of Administration (e.g. 1mg/kg I.V.) | Volume |
|---|---|---|---|
| Tribromoethanol | PBS | 400 mg/kg IP | 0.2 ml/10g |
| Isoflurane | Oxygen | By inhalation to effect:≤5% for induction; 1-2% maintenance | 1-5%@ 1.5L/min |

#### Post Procedure Care

After testing, mice are placed in a recovery box. Half of the box is warmed to 80-86°F and animal has the opportunity to move away from the heat source. To avoid overheating, the temperature inside the cage is monitored with a thermometer at rodent level prior to animal placement, and frequently thereafter. When the animals have fully recovered they are returned to cages and checked once again before the end of the day.

During the time the mouse is recovering from anesthesia, with supportive care as described above, guidelines on supportive care while animal recovers from anesthesia (found in the "Standards for Rodent Survival Surgery at The Jackson Laboratory") will be followed.

#### References (if applicable)

Neal, J. *Protocol: In Vivo Bone Densitometry and Body Composition (DEXA).* February 8, 2017. The Jackson Laboratory.
